# Eye versus hand effector-specificity in the parieto-frontal cortex and cerebellum in humans

**DOI:** 10.64898/2026.10.13.759011

**Authors:** Cristina Rubino, Kiana Masoudi, Jason Gallivan, J. Douglas Crawford

**Affiliations:** Centre for Vision Research, York University, Toronto, Canada; Center for Integrative and Applied Neuroscience, York University, Toronto, Canada; Department of Psychology, York University, Toronto, Canada; Department of Kinesiology & Health Sciences, York University, Toronto, Canada; Department of Biology, York University, Toronto, Canada; Centre for Neuroscience Studies, Queen’s University, Toronto, Canada; Department of Psychology, Queen’s University, Toronto, Canada; Department of Biomedical and Molecular Sciences, Queen’s University, Toronto, Canada

**Author notes:** co-first authorship (Cristina Rubino, Kiana Masoudi). **Corresponding Author**: J. Douglas Crawford Centre for Vision Research, Room 0009A LAS, 4700 Keele Street Toronto, Ontario, M3J 1P3 Canada.

**Keywords:** functional magnetic resonance imaging (fMRI), saccades, reaches, motor planning, sensorimotor processes, univariate analysis, multivariate analysis

## Abstract

The degree and distribution of effector-specific representations for saccade and reach movements in the human brain remain unclear, partly because overlapping regional activation and the limited temporal resolution of fMRI complicate the identification of effector-dependent signals. Using a cue-separated event-related fMRI paradigm, we examined cortical and cerebellar activity across three temporally separated phases: *visual target encoding*, *movement planning*, and *movement execution* in 19 adults. We combined conventional univariate analyses with multivariate analyses of spatial activity patterns to characterize overlapping and distinct neural activity associated with each effector. Univariate analyses revealed widespread overlap across parieto-frontal regions, alongside effector-dependent differences in the magnitude and distribution of activation. Reach-related activity was generally more extensive than saccade-related activity, particularly during planning, while execution revealed distinct patterns of effector-preferential activation across cortical and cerebellar regions. Despite overlapping activation, multivariate analyses identified distinguishable spatial activity patterns across cortical and cerebellar regions. These findings confirm that (1) separating sensorimotor phases improves the detection of effector-preferential signals across the brain, and (2) overlapping activation during eye and hand movements does not necessarily imply equivalent neural activity patterns. Instead, multivariate analyses reveal widespread effector-dependent differences, suggesting a greater degree of spatial differentiation between saccade– and reach-related activity across the human cortex and cerebellum than conventual activation analyses alone would indicate.

## INTRODUCTION

Humans rely on eye and arm movements to interact with the world around them. Both non-human primate and human studies have extensively examined the neural regions involved in controlling visually-guided saccades and reaches. In non-human primate studies, distinct signals related to saccades and reaches have been identified along the posterior parietal cortex (PPC) and frontal regions (Bruce & Goldberg, 1985; Barash et al. 1991; Johnson et al., 1996; Snyder et al., 1997; Andersen & Buneo, 2002). Conversely, in humans, separating saccade– and reach-specific signals has proven more challenging, with most functional magnetic resonance imaging (fMRI) studies reporting substantial overlap between the neural regions recruited for these effectors (Astafiev et al., 2003; Levy et al., 2007; Beurze et al., 2009; Filimon et al., 2009). Consequently, the degree of effector specialization in humans remains unclear.

Across the parieto-frontal cortex, many of the same regions are significantly activated for both reach and saccade tasks (Astafiev et al., 2003; Levy et al., 2007; Beurze et al., 2009; Filimon et al., 2009; Vesia and Crawford 2012). Conjunction analyses of saccade and reach planning reveal overlapping activation in bilateral supplementary motor area (SMA), cingulate motor area (CMA), dorsal and ventral premotor cortex (PMd; PMv), anterior and caudal intraparietal sulcus (aIPS; pIPS), and the parieto-occipital sulcus (PO; Beurze et al., 2009). Findings from Cappadocia et al. (2018) extend these observations to movement execution, demonstrating overlapping activation in the left medial intraparietal sulcus (mIPS), right angular gyrus (AG), precuneus (PCu), superior parieto-occipital cortex (SPOC), lateral occipitotemporal cortex (LOTC), and frontal eye fields (FEF)/PMv. Region-of-interest (ROI) analyses have also reported no significant differences in reach– and saccade-related voxel counts within bilateral intraparietal sulcus area 1 (IPS1) and FEF during execution (Levy et al., 2007). **However, overlapping regional activation does not necessarily indicate that the underlying neural activity patterns are equivalent, raising the question of whether these regions engage common or effector-dependent neural mechanisms during saccade and reach control.**

Although these findings suggest substantial overlap in the neural regions recruited for saccades and reach control, direct comparisons of their activity have also revealed effector-dependent differences. For example, during movement planning, prior to executing a movement action, reach-preferential activity is seen across left parietal regions (IPL, aIPS, SPL, PCu, AG, SMG), the dorsal precentral gyrus, and anterior cingulate (Astafiev et al., 2003; Cappadocia et al., 2018). Bilateral reach-preferential activity also exists in the middle occipital gyrus (MOG), mIPS, primary motor cortex (M1), primary somatosensory cortex (S1), PMd, and BA24 (Cappadocia et al., 2018). During movement execution instead, reach-preferential activity shifts, extending across a broad parieto-frontal network, including bilateral mIPS, SMG, S1, M1, PMd, and SMA, with left-lateralized responses in lateral occipital cortex (LOC), aIPS, IPL, FEF, SMA, anterior PCu, superior frontal gyrus, and BA24 (Astafiev et al., 2003; Hagler et al., 2007; Levy et al., 2007; Filimon et al., 2009; Cappadocia et al., 2018). In contrast, saccades show preferential activity in visual areas, including the bilateral occipital pole and left V7, as well as the bilateral inferior central sulcus and right FEF (Levy et al., 2007; Filimon et al., 2009; Cappadocia et al., 2018).

Although previous studies have identified both overlapping and effector-dependent cortical activity during saccade and reach processing, findings across studies remain difficult to reconcile. This may be due to differences in measurement techniques, task designs, and analytical approaches, as outlined below. (1) One possible explanation for these inconsistent results is the temporal overlap of sensorimotor processes, which can be difficult to disentangle using fMRI, given its temporal resolution on the order of seconds. Consequently, signals associated with different brain regions or temporal events that occur close together might “blur” together. This temporal conflation may reduce the detection of activity involved in preparing and executing saccades or reaches (Cappadocia et al., 2017). (2) Many previous studies have employed task designs that do not explicitly separate the three distinct phases involved in performing visually guided movements: perceiving a peripheral target, planning a movement towards its remembered location, executing that movement. Studies that present a target and instruct an effector (i.e., hand or eyes) during the same phase might produce results that reflect both visual target encoding and motor planning. Moreover, studies that do not separate planning and execution phases may show effector-independent activation in brain regions that contribute to saccades and reaches at different time points. For example, Cappadocia et al., (2017; 2018) used an event-related task that distinguished these three phases, revealing a distinct temporal progression of cortical sensorimotor activity and a medial-lateral organization of effector-preferential activation. (3) Additionally, traditional univariate analyses limit the types of information that can be extracted from brain regions. While saccade and reach regions may exhibit overlapping activation, and direct contrasts may reveal evidence of effector-specificity, these approaches are all based on the magnitude of the blood-oxygen-level-dependent (BOLD) signal and rather than finer-grained information contained in spatial activity patterns across voxels. Previous multivoxel pattern analyses demonstrated that effector-dependent and effector-independent movement information can coexist within individual parieto-frontal regions, even in the absence of differences in mean activation (Gallivan et al., 2011), highlighting how different analytical approaches can reveal nuances in neural activity patterns that may not be captured by conventional univariate analyses.

Further, overlapping activation at the regional level does not necessarily establish that saccades and reaches elicit similar underlying activity patterns (Kriedeskorte et al., 2006; Diedrichsen & Kriegeskote 2017). Likewise, an absence of differences in mean activation does not preclude effector-dependent differences in the spatial distribution of activity within a region (Kriedeskorte et al., 2006). Thus, it remains unclear how broadly distinguishable multivoxel activity patterns occur within regions exhibiting overlapping activation or comparable mean activity. Combining temporally separated sensorimotor phases with multivariate pattern dissimilarity analyses may therefore provide a more comprehensive characterization of effector-dependent neural activity beyond conventional univariate comparisons.

An additional gap concerns the cerebellum, where the spatial organization of effector-dependent activity remains less well characterized than in the cortex. Few fMRI studies have examined cerebellar activation overlaps and differences between saccades and reaches, and they typically focus on separate task contrasts and qualitative comparison of the resulting activity rather than more direct approaches within the same experimental paradigm (Miall et al., 2000; Nitschke et al., 2005).

The present study employed a cue-separation fMRI paradigm (Cappadocia et al., 2018) to better understand effector-dependent activation across temporal phases of sensorimotor control within cortical and cerebellar regions. Specifically, we examined how functional activity within the saccade and reach networks: (1) varies across *visual target encoding*, *movement planning*, and *execution*; (2) which regions exhibit overlapping activation between effectors; (3) and which regions are preferentially activated for saccades versus reaches during planning and execution. We further investigated (4) whether saccades and reaches elicit reliably distinguishable multivoxel activity patterns within cortical and cerebellar regions.

The first three questions were addressed using conventional univariate analyses, through whole-brain voxel-wise analyses of the spatial *distribution* of activation and ROI analyses of action *magnitude* within established saccade– and reach-related regions. The fourth question was examined using a multivariate approach involving the calculation of crossvalidated squared Euclidean distances between saccade– and reach-related multivoxel activity patterns within ROIs.

We hypothesized that (1) shared, effector-independent cortical activity would occur during visual target encoding, with saccade– and reach-related cortical (parieto-frontal) and cerebellar activity emerging during *movement planning* and *execution*. Reach planning and execution were expected to show broader activity, lateralized to the left hemisphere (i.e., contralateral to the reaching hand). (2) Overlapping activation between saccades and reaches was expected across key cortical (parieto-frontal) and cerebellar regions, with greater overlap during *movement execution*. (3) During *movement planning*, reach-preferential cortical (parieto-frontal) activity was expected to be more widespread compared to saccades. During *movement execution*, we predicted a relative medial-lateral separation of reach– and saccade-preferential activation, respectively, with reach-preferential activity more left-lateralized. Within the cerebellum, saccade-preferential activity was expected in medial regions (i.e., oculomotor vermis), while reach-preferential activity was expected in the ipsilateral cerebellar hemisphere. (4) Despite overlapping cortical and cerebellar activation during movement planning and execution, we hypothesized that saccades and reaches would elicit distinguishable multivoxel activity patterns within these regions.

## METHODS

### Participants

Twenty, right-handed, healthy adults, with normal or corrected-to-normal vision were recruited from local communities, predominantly from York University. An a priori power analysis was conducted using effect magnitudes in similar visuomotor studies, which observed approximately 0.5% to 1.5% blood-oxygen-level-dependent (BOLD) signal change in pertinent regions of interest across the cortex (Gallivan et al. 2011; Cappadocia et al. 2017). Assuming a large effect size (Cohen’s d = 0.8-1.00), ɑ = 0.05, and power (1-β) = 0.80, the analysis indicated 10 to 15 participants would be sufficient to detect expected effects. To account for exclusion of participants due to potential motion artifacts or behavioural errors, an additional five participants were recruited, resulting in the final sample of 20. The York University Human Participants Review Sub-committee approved all methods used in the study. All participants provided informed, written consent prior to the start of the experiment.

### Experimental Apparatus

The experimental apparatus of the present study, shown in **Figure 1**, was similar to Chen et al. (2014) and Cappadocia et al. (2017). Visual stimuli consisted of light emitting diodes (LEDs) that were projected onto a custom-built screen board via fixed optic fibers. A translucent touch screen (Keytec, 170 x 126 mm) was attached to the top of the screen to record touch responses. The screen was fixed to a custom-built platform that was placed over the participant’s abdomen in the MRI, with its exact distance adjusted to ensure comfortable reaching. An SR Research EyeLink 1000 Plus eye-tracker was attached to the MRI bed using an adjustable holder, to the side and behind the platform. The exact position of the camera was modified for clear viewing of the participant’s left eye. Additionally, five independent custom-built illuminators were attached to the platform using adjustable holders, positioned above the screen to illuminate the eye.

**Figure 1:**
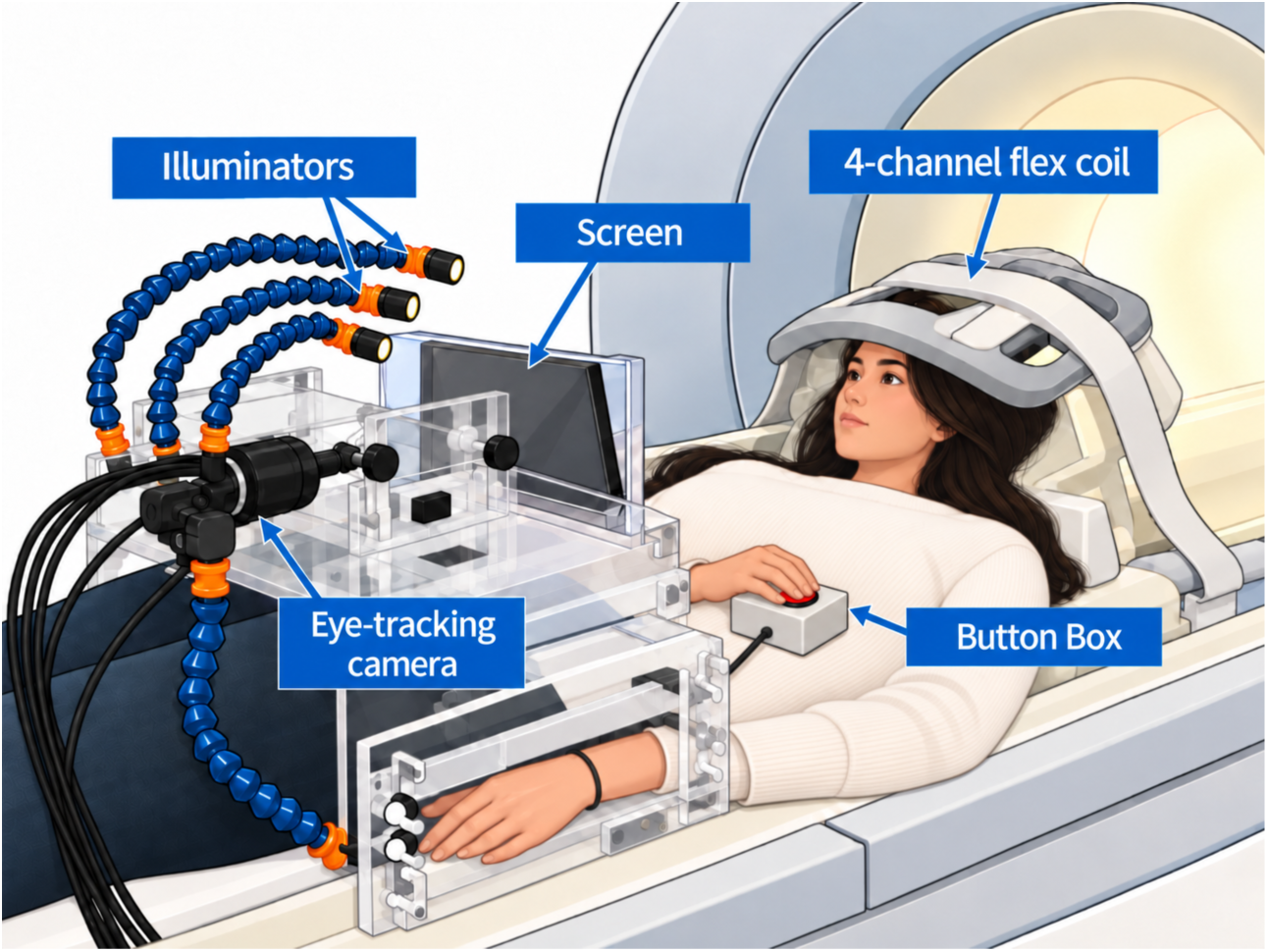
Image of the experimental set-up. Participants performed the experimental tasks in the MRI bore by interacting with a screen positioned above their torso. A button box placed on their abdomen was used for the right hand’s ‘home’ or ‘starting’ position and for responses during the control task. Left eye-movements were recorded using an eye-tracking camera, with five illuminators mounted above the screen to provide illumination for the eye.

The participant’s head was tilted ∼20° to allow for a comfortable view of the screen. The left arm rested alongside the body while the right arm was bent to allow the hand to rest on a button box placed on the abdomen. This marked the ‘home’ or ‘starting’ position. Participants wore MRI-compatible headphones to hear task-related instructions and communicate with researchers between experimental runs. Throughout the entire imaging session, the MRI room was completely dark, with the only light coming from the LEDs on the touch screen and the eye-tracking illuminators. As such, participants were not able to see their hand during trials, up until the movement execution phase, at which point it was possibly visible as the hand got closer to the screen.

### Experimental Paradigm

An event-related design was used, with the paradigm including three tasks: saccade (“look”), reach (“reach”), as well as colour-report (“colour”), which served as the control. Participants completed these tasks with the support of visual stimuli presented on the screen. These consisted of one yellow central fixation dot and red or green peripheral targets that were presented 4-7° left or right of the midline, on the same horizontal plane. In addition, twenty white “mask” LEDs were presented during delay phases of each trial, with ten appearing on each side of the midline, five below and five above the horizontal target line. These masks were used to control for visual after images of the visual targets and encouraged participants to focus on remembered stimuli (Sereno et al. 2001; Medendorp et al. 2003). All visual stimuli had a diameter of 3mm. Information regarding the stage at which these visual stimuli appeared within a trial is outlined in the following section.

**Figure 2** illustrates the sequence of events in each trial. Each run of trials began with 12 seconds of central fixation before the start of the first trial to help stabilize the BOLD signal. Subsequent trials were cued with the flashing of the central yellow LED at the end of a preceding trial.

**Figure 2:**
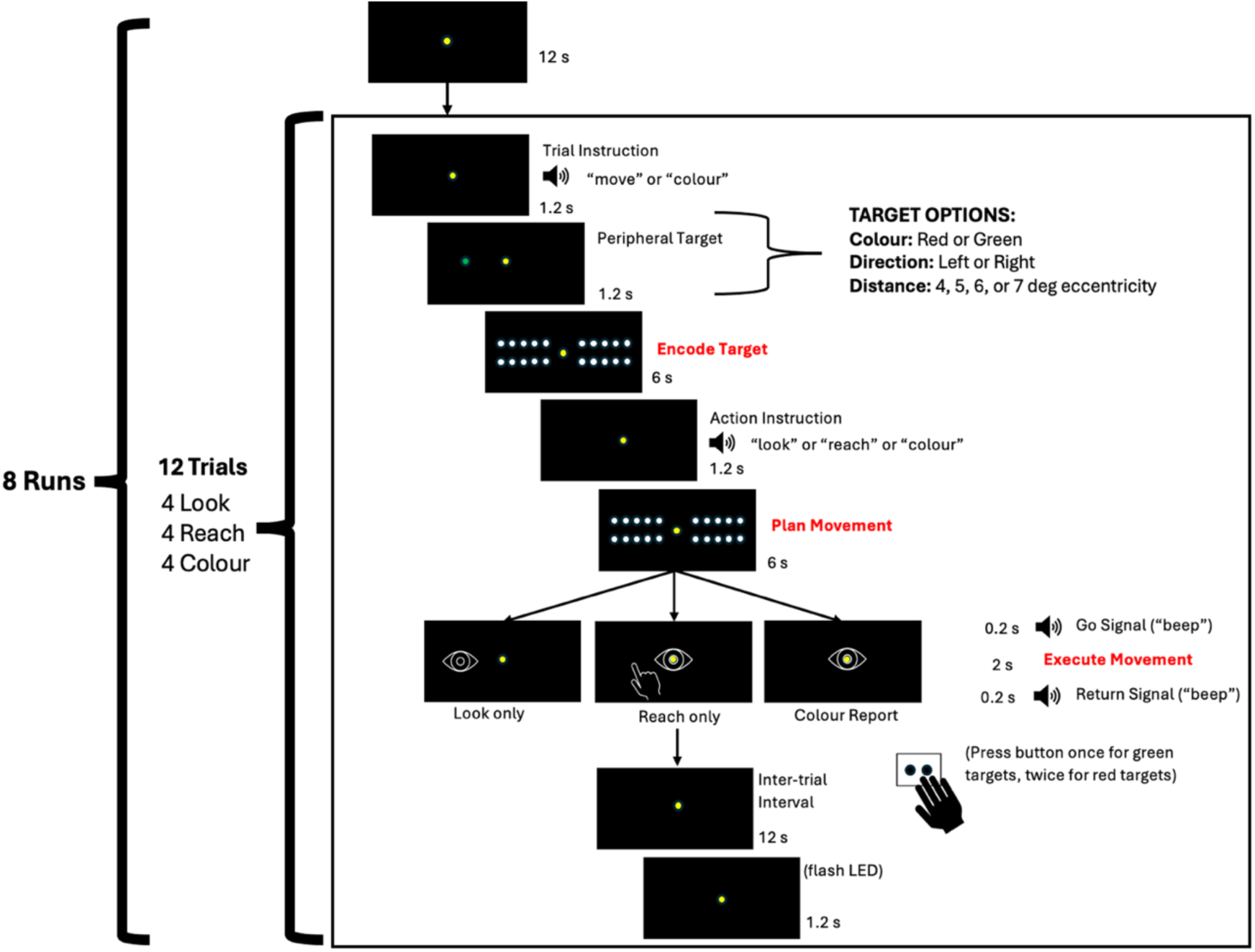
Illustration of the experimental paradigm. Visual stimuli on black screens represent participants’ view within each timeframe of a single trial. The difference between the two experimental conditions (Saccade and Reach) emerged from the auditory instructions, which dictated the movements participants performed towards the remembered target position. The control condition (colour-report) required participants to remember the colour of the peripheral target and report it using the button box.

Participants were instructed to keep their hand and eyes at the ‘home’ position (i.e., their right hand on the button box and eyes fixating the central yellow LED) through the entire trial until the *movement execution* phase. During the first 1.2 seconds, participants were told if this was a “move” or “colour” trial. A red or green peripheral target then appeared either on the left or right side of the yellow LED. Following this, the *visual target encoding* phase began, with white mask LEDs presented for 6 seconds while participants continued fixating. During the 1.2 second period after the mask disappeared, an auditory instruction indicated either “reach”, “look”, or “colour” (initiating the effector-dependent planning phase). This was followed by the second 6-second delay period, corresponding to the *movement planning* phase, during which time the white LED masks reappeared. Following this, participants heard a “beep” sound, which served as the go-signal to initiate the *movement execution* phase. In the saccade trials, participants shifted their gaze to fixate towards the remembered target location until hearing a second “beep”, which prompted a return to central fixation. In the reach trials, participants touched the board with their right index finger at the remembered target position while maintaining central fixation, holding that position until the return signal prompted them to move their right arm back to the button box. Finally, in the colour condition, participants maintained central fixation while indicating target colour by pressing the home button with their right index finger, once for green and twice for red. The response window of the movement execution phase was 2.4 seconds, after which a 12-second intertrial interval began. Each complete trial was 31.2 seconds.

Each functional run consisted of 12 trials (four per condition). Different trial types (saccade, reach, colour-report) were presented in a random order, with 50% of peripheral targets presented in each visual hemifield for each condition. No two trials repeated the same identical condition. Each session consisted of eight functional runs inside the scanner, with a break between the 4^th^ and 5^th^ run.

### Training and Behavioural Recordings

Participants attended a training session at the MRI Simulator Facility at the Sherman Health Sciences Centre at York University at least 24 hours prior to the scan, which lasted approximately 30 minutes. During this time, we first confirmed with participants whether they were able to remain in the scanner for the entire duration of the experiment. We then explained the task and had them complete a few practice trials using the experimental equipment until they were confident they understood the task. In addition to this, prior to scanning participants on the day of, they completed an additional run outside of the MRI bore, referred to as “Run 0.” This allowed us to ensure that participants performed the task correctly and have a record of consistent behaviour in cases where eye-tracking was lost inside the scanner. Prior to Run 0, the position of the eye-tracker was adjusted to allow for viewing of the left-eye. Pupil and corneal reflection thresholds were adjusted on the EyeLink host computer, and a custom calibration task was employed to capture the x and y position of the pupil on the board. During calibration, participants sequentially fixated on 7 LEDs for 2 seconds each, with each position shown twice, achieving an average accuracy of 0.25-0.5 degrees (SR Research Ltd., 2025).

### Imaging Parameters

The scanning sessions took place at the York MRI Facility at the Sherman Health Sciences Centre at York University with a 3-T Siemens Magnetom TIM Trio scanner. The posterior section of a 12-channel head coil (6 channels) was positioned behind the participant’s head and 4-channel flex coil over the front.

Functional images were collected using an echo-planar imaging (EPI) sequence (repetition time (TR) = 1200 ms; echo time (TE) = 30 ms; flip angle (FA) = 66°; field of view (FoV) = 240 mm x 240 mm; matrix size = 96 x 96, producing an in-plane resolution of 2.5 x 2.5 mm; slice thickness = 3 mm; no gap; 54 transverse slices captured in an interleaved order). Acquisition time was 9 minutes and 22 seconds.

Structural, T1-weighted (T1-w) images were acquired using an MPRAGE sequence (TR = 2300; TE = 2.26 ms; inversion time (TI) = 900 ms; FA = 8°; FoV = 256 x 256; matrix size = 256 x 256, producing an in-plane resolution of 1.0 mm x 1.0 mm; slice thickness = 1.0 mm; no gap; 192 sagittal slices captured in an ascending order) for whole brain structural imaging. Acquisition time was 5 minutes and 21 seconds.

To correct for distortion, dual-echo gradient field maps were acquired using a GRE sequence (TR = 566 ms; TE1 = 4.92 ms; TE2 = 7.38 ms; FA = 60°; FoV = 240 mm x 240 mm; matrix size = 96 x 96, producing an in-plane resolution of 2.5 x 2.5 mm; slice thickness = 3mm; no gap; 54 transverse slices captured in an interleaved order). Acquisition time was 1 minute and 51 seconds.

### Preprocessing

Data preprocessing was conducted using fMRIPrep (Version 24.1.1; Esteban et al. 2018; 2019) and FMRIB Software Library (FSL; Version 6.00; Jenkinson et al., 2012). T1-w scans were skull-stripped using deepbet (Fisch et al., 2024) before participant-level pre-processing on all functional and anatomical data was executed using fMRIPrep. fMRIPrep performed minimal preprocessing, including non-uniformity correction, brain masking of the T1-w image, tissue segmentation, head-motion correction of the functional images, susceptibility distortion correction using acquired field-maps, co-registration of each functional run to the T1-w image using boundary-based registration, and nonlinear spatial normalization of T1-w (and consequently the functional scans) to standard MNI template space (MNI152Lin6Asym).

Following fMRIPrep, each preprocessed BOLD run was skull-stripped prior to running additional preprocessing and processing in FSL’s FEAT. Additional pre-processing in FSL during first-level analysis included spatial smoothing (8 mm Full Width at Half Maximum (FWHM)) and high-pass temporal filtering (100 s).

### Data Analysis

#### Behavioural Analysis and Exclusion Criteria

To ensure participants correctly performed the instructed tasks during each trial, the event timing files for functional runs were examined offline. A trial was excluded from analysis if: 1) an uninstructed reach behaviour was recorded (e.g., participants touched the board for a colour-report trial) and 2) no action was recorded (e.g., participants did not reach to touch the board). In four participants, equipment issues prevented touch-end points from being recorded, despite participants reaching and touching the board with their index finger. In such cases, we verbally confirmed with participants that they had indeed touched the board for reach trials following each run, although we were not able to verify behaviour further from this. Reach trials were not excluded for these scanning sessions. Additionally, the eye was monitored in real-time from the EyeLink Host monitor during the scan to confirm that participants were awake and performing the task. Although the eye-tracking camera was active during the scanning session, the setup did not provide optimal illumination and positioning for complete gaze tracking data during scans for any participant. Thus, we could not remove trials based on this data. Instead, eye-tracking data from Run 0 served as confirmation that participants were able to comply with the behavioural requirements of the task (i.e., fixate during trials and shift gaze during saccade execution). The touchscreen recorded time and location of the reach end point.

Trials were also excluded if framewise displacement (FD) exceeded 0.99 mm inside the scanner outside the execution and inter-trial interval (ITI) phases. Runs were excluded if more than five trials, or more than two from the same condition, were removed due to head motion.

Due to technical malfunctions, four participants did not complete the full eight function runs with one to four runs missing per participant, resulting in a total of 10 missing runs. Following preprocessing, one participant was excluded due to excessive head motion (all eight runs; 128 trials), leaving 19 participants (142 runs; 1704 trials) for final analysis. Of these, 32 trials (1.88%), including two entire runs, were removed due to excessive motion, with a mean of 1.68±4.32 trials per participant. Additionally, 43 trials (2.52%) were removed due to behavioural errors, with a mean of 2.26±1.66 trials per participant. Maximum number of trials excluded for a participant included in the analysis was five. In total, 75 trials (4.40%) were excluded across 19 participants, with a mean of 3.95±4.47 trials per participant, leaving 140 runs and 1629 trials for final analysis.

#### Univariate Analysis

Functional data was analyzed using FSL (Version 6.00; Woolrich et al., 2001). Peripheral target locations were pooled across left and right directions as laterality was not a factor of interest in the following analyses. Functional runs were fitted to three separate general linear models (GLMs) to examine task-related activation during 1) *visual target encoding* 2) *movement planning* and 3) *movement execution* phases. For the *visual target encoding* phase, two predictors were defined based on condition: *Move* and *Colour.* One contrast was created to examine activation for encoding the spatial location of target for future movement versus encoding non-spatial properties for reporting colour: Contrast 1 [Move>Colour]. For the *movement planning* phase, three predictors were defined based on condition: *Reach*, *Saccade*, and *Colour*. Two contrasts were designed to investigate effector-related activation during saccade and reach planning compared to colour-report (control): Contrast 2 [Saccade>Colour] and Contrast 3 [Reach>Colour], and additional two contrasts were defined to investigate regions with greater activity for one effector over the other (effector-specificity contrasts): Contrast 4 [Saccade>Reach] and Contrast 5 [Reach>Saccade]. An identical GLM was set up for the *movement execution* phase (Contrasts 6 to 9). White matter time series (derived from fMRIPrep) was included as a nuisance regressor.

Parameter estimates were combined across runs within each participant using a fixed-effects model. Group-level statistical inference was performed using FSL randomise (Winkler et al., 2014), which performed voxel-wise non-parametric permutation testing (5000 permutations) with threshold-free cluster enhancement (TFCE; Smith et al., 2009) and family-wise error (FWE) correction (p<0.05). The *cluster* function on FSL was used to report cluster information for each contrast. Group level thresholded statistical maps were projected from volumetric MNI152 space onto FreeSurfer fsaverage surface using mri_vol2surf registration for visualization.

The anatomical location of cortical peak voxels and relevant local maxima were labelled using probabilistic atlases in volumetric space: the Harvard-Oxford Cortical Structural Atlas and Harvard-Oxford Subcortical Structural Atlas, provided by the Harvard Centre for Morphometric Analysis. The probability values reflect the proportion of individuals in the atlas sample for whom a given voxel was assigned to a particular region (FMRIB Software Library, n.d.). Cerebellar areas were identified using the Cerebellar Atlas in MNI152 space after normalization with FNIRT. All peak voxels were labelled unless they were too small (i.e., less than 10 voxels) or predominantly in white matter regions. Additionally, some local maxima within clusters were labelled to provide further information on the extent of regions activated for each contrast. To estimate the location of MNI coordinates on the fsaverage inflated brain, a publicly available Yeo Lab function was used, which performs precomputed volumetric to surface mapping and outputs corresponding surface vertex values (Wu et al., 2018). Since the anatomical labels were derived from volume space, their spatial location appeared slightly shifted in some cases on the cortical surfaces.

#### Conjunction Analysis

To identify regions that are activated for both reach and saccade during the *movement planning* and *movement execution* phases, binary masks of the TFCE-corrected, t-statistic maps (thresholded at FWE p<0.05) for [Saccade>Colour] and [Reach>Colour] contrasts were generated. These binary maps were multiplied voxel-wise using *fslmaths*, to produce a conjunction map of overlapping areas.

#### Region of Interest Analysis

To further determine activity across key regions in the parieto-frontal cortex, ROIs were selected using coordinates reported in previous literature. These included: bilateral anterior cingulate cortex (aCing); bilateral dorsolateral prefrontal cortex (DLPFC); supplementary eye field (MedFG); bilateral frontal eye field (MidFG); bilateral M1; bilateral PMd; bilateral AG; bilateral SPOC; bilateral mIPS; bilateral S1; bilateral primary visual cortex (V1). The MNI coordinates and corresponding references used for these regions is shown in **Table 1**.

**Table 1:** Name of regions selected for ROI analysis, hemisphere, MNI coordinates, and corresponding references.

| **Region** | **Hemisphere** | **MNI**<br>**X** | **MNI**<br>**Y** | **MNI**<br>**Z** | **References** |
| --- | --- | --- | --- | --- | --- |
| aCing | Left | -4 | 16 | 32 | Jamadar et al., (2013) |
| aCing | Right | 6 | 8 | 38 | Jamadar et al., (2013) |
| DLPFC | Left | -38 | 34 | 36 | Cameron et al., (2015) |
| DLPFC | Right | 32 | 40 | 34 | Jamadar et al., (2013) |
| MedFG |  | -2 | -10 | 64 | Jamadar et al., (2013) |
| MidFG | Left | -40 | -14 | 54 | Jamadar et al., (2013) |
| MidFG | Right | 40 | -8 | 54 | Jamadar et al., (2013) |
| M1 | Left | -39 | -23 | 59 | Sartin et al., (2023) |
| M1 | Right | 35 | -25 | 64 | Sartin et al., (2023) |
| PMd | Left | -25 | -10 | 61 | Cappadocia et al., (2017) |
| PMd | Right | 22 | -11 | 64 | Cappadocia et al., (2017) |
| AG | Left | -40 | -70 | 44 | Vesia et al., (2010) |
| AG | Right | 35 | -75 | 46 | Vesia et al., (2010) |
| SPOC | Left | -10 | -85 | 46 | Vesia et al., (2010) |
| SPOC | Right | 9 | -85 | 45 | Vesia et al., (2010) |
| mIPS | Left | -22 | -68 | 45 | Vesia et al., (2010) |
| mIPS | Right | 22 | -66 | 45 | Vesia et a., (2010) |
| S1 | Left | -40 | -28 | 52 | Mayka et al., (2006) |
|  |  |  |  |  | Blohm et al., (2019) |
| S1 | Right | 38 | -30 | 42 | Mayka et al., (2006) |
|  |  |  |  |  | Blohm et al., (2019) |
| V1 | Left | -6 | -80 | 10 | Desouza et al., (2003) |
| V1 | Right | 9 | -79 | 8 | Desouza et al., (2003) |
| Lobule VI | Left | -13 | -65 | -25 | Simo et al. (2005) |
| Lobule VI | Right | 28 | -54 | -28 | Errante et al. (2021) |
| Vermis VI |  | -1 | -64 | -36 | Simo et al. (2005) |
| Lobule VIIIb | Left | -20 | -54 | -50 | Errante & Fogassi (2020) |
| Lobule VIIIb | Right | 18 | -62 | -50 | Errante & Fogassi (2020) |
| Crus I | Left | -30 | -68 | -45 | Simo et al. (2005) |
| Crus I | Right | 30 | -76 | -38 | Simo et al. (2005) |

Spherical ROIs (6-mm radius) were created at the centre of each coordinate using FSL to generate binary masks for each region. For each participant and contrast, ROIs were restricted to voxels within the subject-specific brain mask generated during higher-level FEAT analysis. ROI statistics were extracted for [Saccade>Colour] and [Reach>Colour] contrasts (Contrasts 2-3 and 6-7) using *featquery*, which computed percent signal change for contrast of parameter estimates (COPEs) within each masked ROI.

#### Multivariate Analysis

##### First-Level GLM

To estimate condition-specific activity patterns for multivariate analyses, a separate first-level GLM was implemented in FSL FEAT for each functional run. The model included separate regressors for all task-related conditions (i.e., instructions, peripheral target), including separate regressors for saccade, reach, and colour-control conditions during planning and execution. Task regressors were convolved with the hemodynamic response function, and temporal derivatives were omitted. Temporal autocorrelation was addressed using FEAT’s FILM prewhitening procedure. No spatial smoothing was applied (0-mm FWHM) to preserve fine-grained multivoxel activity patterns. All other preprocessing procedures and model parameters were preserved from the univariate analyses, except for the modifications described above.

Individual contrasts were specified to estimate activity associated with each experimental condition relative to the implicit baseline. The resulting condition-specific parameter estimate images were extracted from each ROI and functional run for subsequent multivariate analyses.

To determine whether saccade and reach conditions elicited distinguishable multivoxel activity patterns within overlapping cortical and cerebellar regions, we calculated crossvalidated squared Euclidean distances (Kriegeskorte et al., 2008; Walther et al., 2016) between their voxelwise parameter estimates within each ROI, separately for planning and execution.

For each participant, ROI, and run, a difference pattern was calculated by subtracting the reach parameter estimates from the corresponding saccade estimates. Crossvalidation was performed by calculating the inner product of these difference patterns across every unique pair of independent runs, normalized by the number of voxels within the ROI, and averaging the resulting distances to obtain one estimate per participant and ROI:

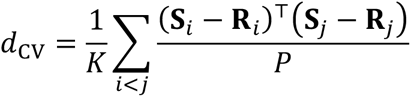

where (S) and (R) represent the voxelwise saccade and reach parameter estimate vectors, respectively, (P) is the number of voxels within the ROI, and (K) is the number of unique run pairs. By comparing difference patterns across independent runs, crossvalidation reduces positive noise bias associated with within-run distance estimates, allowing distances to take negative values due to measurement variability.

Two complementary analyses were performed. One used raw parameter estimates, preserving differences in both overall response magnitude and spatial activity patterns. A second analysis removed the mean voxel response separately for each condition, run, and ROI before calculating distances, isolating differences in relative spatial activity patterns independent of uniform shifts in regional activation magnitude. Specifically, voxel demeaning was performed as follows:

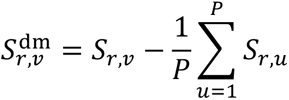

where (r) denotes the run, (v) the voxel, and (P) the number of voxels within the ROI. The demeaned saccade and reach patterns were then entered into the same crossvalidated distance calculation described above. No variance scaling was applied following mean removal, preserving the magnitude of voxelwise deviations from the regional mean.

#### Statistical Analysis

##### Univariate Analyses

Statistical tests were performed using R (Version 4.5.3). To determine whether activation significantly differed from the colour condition, mean percent signal change for each contrast within each region and phase was assessed using one-sample, two-sided t-tests. P-values were corrected using false discovery rate (FDR) to control for multiple comparisons. To directly compare activation between effectors (saccades [Saccade>Colour] versus reaches [Reach>Colour]), percent signal change for each region was assessed using paired t-tests. P-values were corrected using FDR within each phase.

##### Multivariate Analysis

At the group level, distances were evaluated using two-sided one-sample t-tests against zero, separately for each ROI, phase, and analysis. Statistical significance was assessed using Benjamini–Hochberg false discovery rate correction across 28 ROIs (q = 0.05), independently for planning and execution and for raw and mean-removed estimates. Group means and 95% confidence intervals were calculated across participants. Significantly positive group-level crossvalidated distances were interpreted as evidence of reliably distinguishable saccade and reach multivoxel activity patterns within an ROI, rather than differences arising solely from positive within-run noise bias.

## RESULTS

The present study examined four questions concerning saccade– and reach-related activity across cortical and cerebellar regions: (1) the distribution of activation across visual target encoding, movement planning, and execution; (2) the extent of overlapping activation between effectors; (3) regional effector preferences; and (4) the distinguishability of saccade– and reach-related multivoxel activity patterns. We hypothesized that visual target encoding would elicit shared activity, whereas planning and execution would reveal effector-dependent activity. Reach-preferential activity was expected in medial parieto-frontal regions and ipsilateral cerebellum, and saccade-preferential activity in lateral parietal and medial cerebellar regions. Finally, despite overlapping regional activation, we predicted distinguishable multivoxel activity patterns between effectors during planning and execution.

To address these questions, we first conducted univariate analyses, examining task-related activation relative to the control condition, overlapping activation through conjunction analyses and effector-preferential differences through direct task contrasts, and activation magnitude differences within predefined ROIs. We then conducted multivariate analyses using crossvalidated pattern distances.

### Effector versus colour activation across encoding, planning, and execution

The first objective of this study was to characterize saccade and reach activation across the *visual target encoding*, *movement planning*, and *movement execution* phases. Contrast 1 ([Move>Colour]) was used for the *visual target encoding* phase and contrasts 2, 3, 6, and 7 ([Saccade>Colour] and [Reach>Colour]) were used for the effector-dependent phases. **Table 2** provides the list of abbreviations used in the labelling cortical regions, as well as references for previous saccade and reach-fMRI studies that have reported activation in these areas.

**Table 2:**
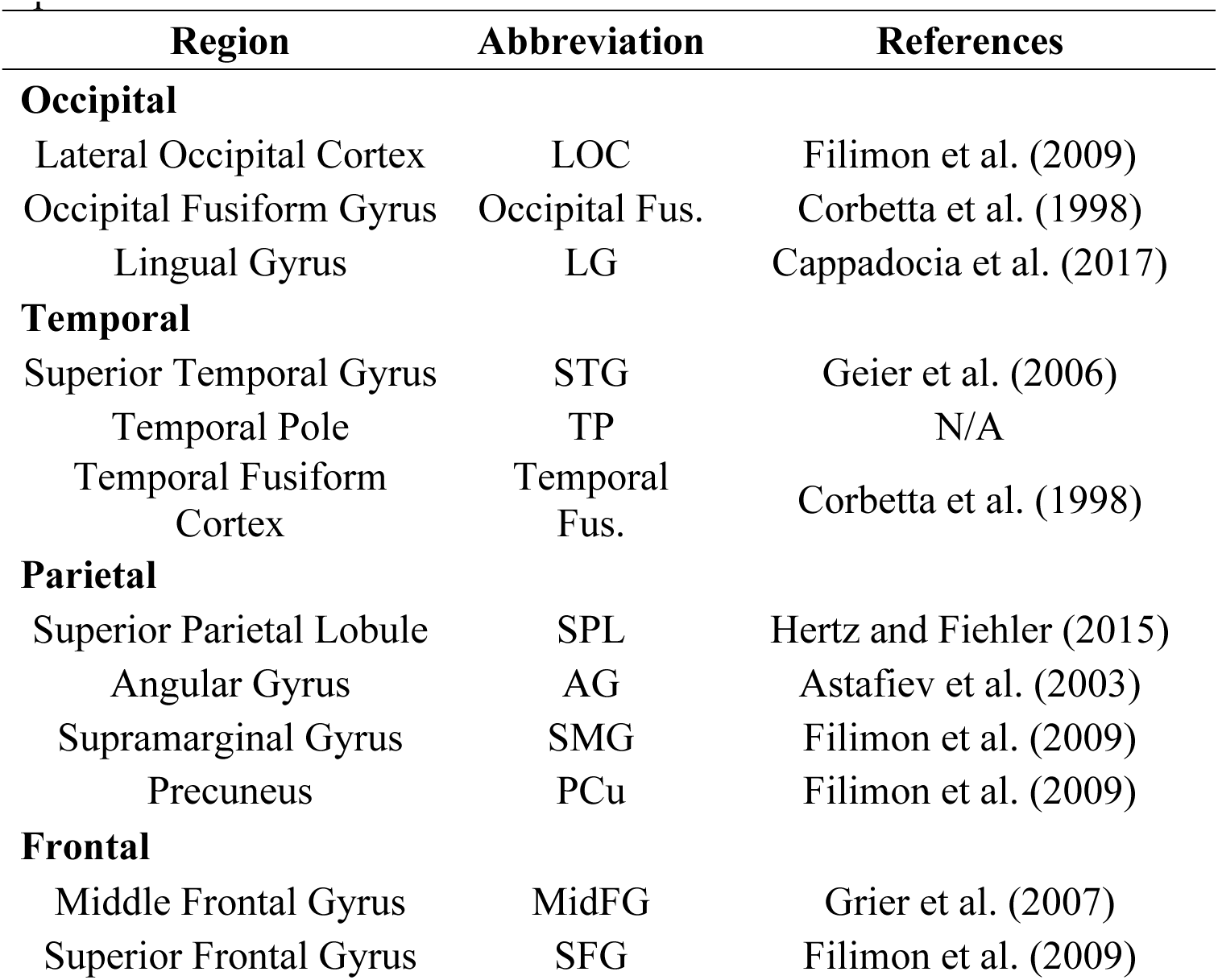
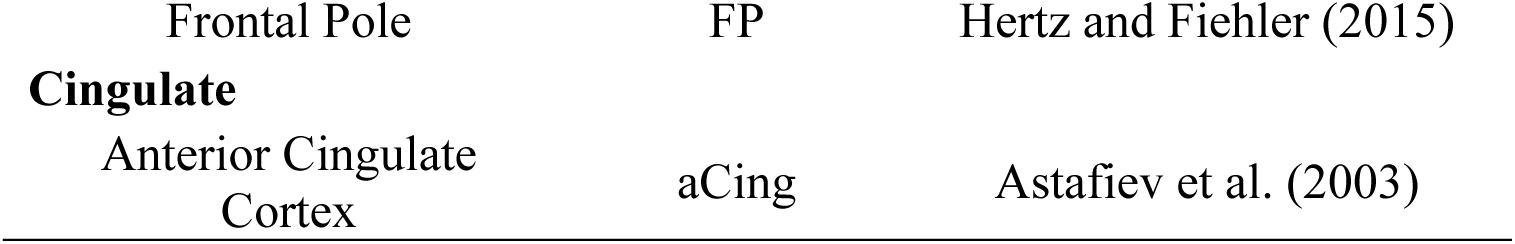
List of brain region abbreviations and saccade or reach-related fMRI literature that have reported activation in these areas.

#### Visual Target Encoding

During the *visual target encoding* phase, participants were instructed on whether they needed to remember the position of the target to perform a movement towards it, or its colour to report. Since there were no instructions on if the movement would require the hand or the eyes, clusters surviving the [Move>Colour] contrast could be related to spatial-working memory or effector-independent movement preparation. **Figure 3** shows the functional map of significant regions, overlaid on the fsaverage inflated cortical surface. **Table 3** reports the MNI coordinates and t-statistic values of peak voxels from significant clusters. During this phase, activation was limited to the left hemisphere, with peak voxels in the TP, STG, and anterior and posterior MidFG.

**Figure 3:**
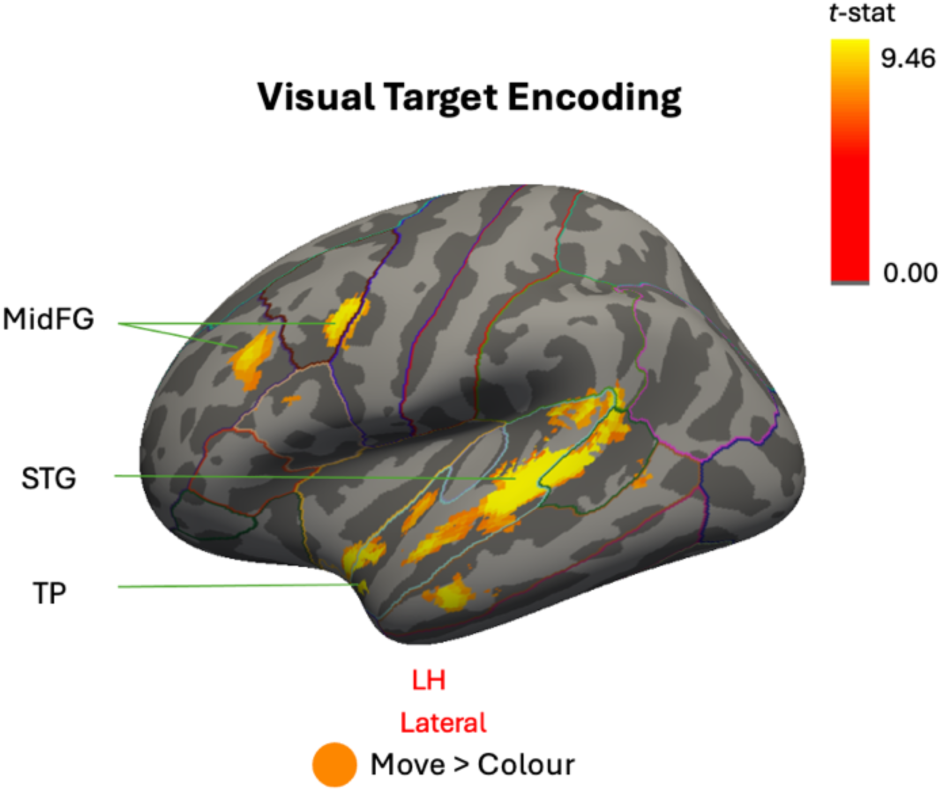
Group-level activation for the visual target encoding phase ([Move>Colour] contrast) overlaid on the FreeSurfer fsaverage ‘inflated’ cortical surface. Light gray represents gyri and dark gray represents sulci. Lines on the surface denote the anatomical boundaries from the Desikan-Killiany Atlas, provided for visualization. Highlighted regions represent significant voxels from TFCE-corrected t-statistic maps (FWE p<0.05), with the colour-gradient corresponding to the magnitude of the t-statistic, as indicated by the colour bar. Labels are peak voxels.

**Table 3:**
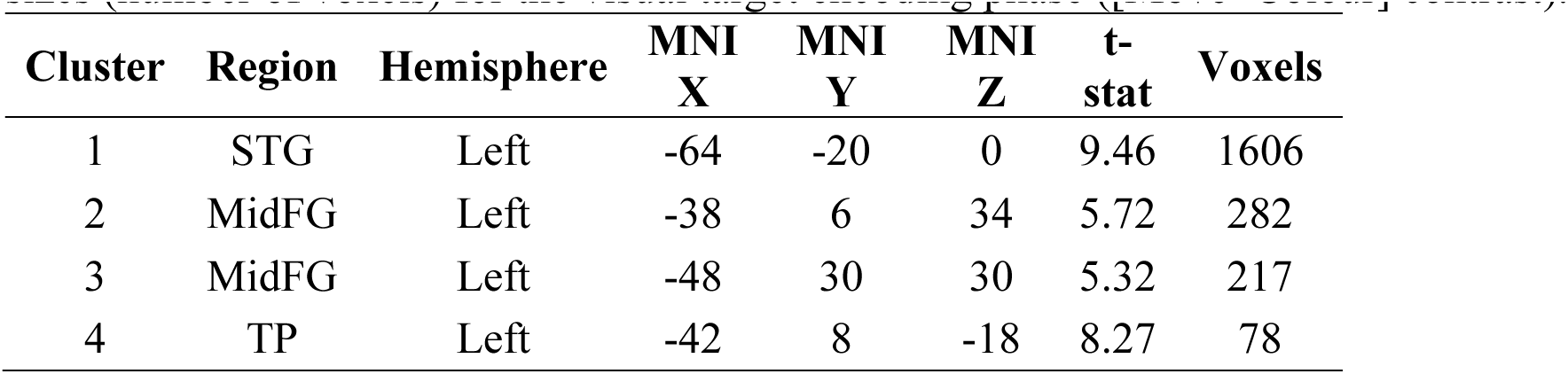
Name of region, hemisphere, MNI coordinates, t-statistic at the coordinate, and cluster sizes (number of voxels) for the visual target encoding phase ([Move>Colour] contrast).

| Cluster | Region | Hemisphere | MNI<br>X | MNI<br>Y | MNI<br>Z | t-stat | Voxels |
| --- | --- | --- | --- | --- | --- | --- | --- |
| 1 | STG | Left | -64 | -20 | 0 | 9.46 | 1606 |
| 2 | MidFG | Left | -38 | 6 | 34 | 5.72 | 282 |
| 3 | MidFG | Left | -48 | 30 | 30 | 5.32 | 217 |
| 4 | TP | Left | -42 | 8 | -18 | 8.27 | 78 |

In summary, greater activation during the *visual target encoding* phase for memory-guided movements was modest and limited to left temporal and middle frontal regions relative to the non-spatial task.

#### Movement Planning

During the *movement planning* phase, participants were aware of the required effector (eye vs. hand). Thus, the contrasts [Saccade>Colour] and [Reach>Colour] isolate effector-dependent planning activity relative to preparing for a button-press (colour-report condition). **Figure 4** shows these cortical functional maps, overlaid on the fsaverage inflated cortical surface. **Table 4** reports the MNI coordinates and t-statistic values of peak voxels, along with select local maxima within each significant cluster to illustrate the spatial extend of activation. Saccade planning produced activation primarily across posterior parietal regions (**Figure 4A**). The largest cluster (1041 voxels) revealed peak and local maximum activity in the left anterior and posterior SPL. A smaller homologous cluster (262 voxels) was observed in the right hemisphere, with peak activity in right SPL and local maximum within PCu. Additionally, a smaller cluster (22 voxels) was present in the left precentral gyrus. Conversely, reach planning was characterized by a single, large cluster (80445 voxels), spanning the occipital-parietal-frontal cortex (**Figure 4B**). The peak voxel was located in the left SPL, and local maxima were present in frontal regions, specifically the SFG and FP.

**Figure 4:**
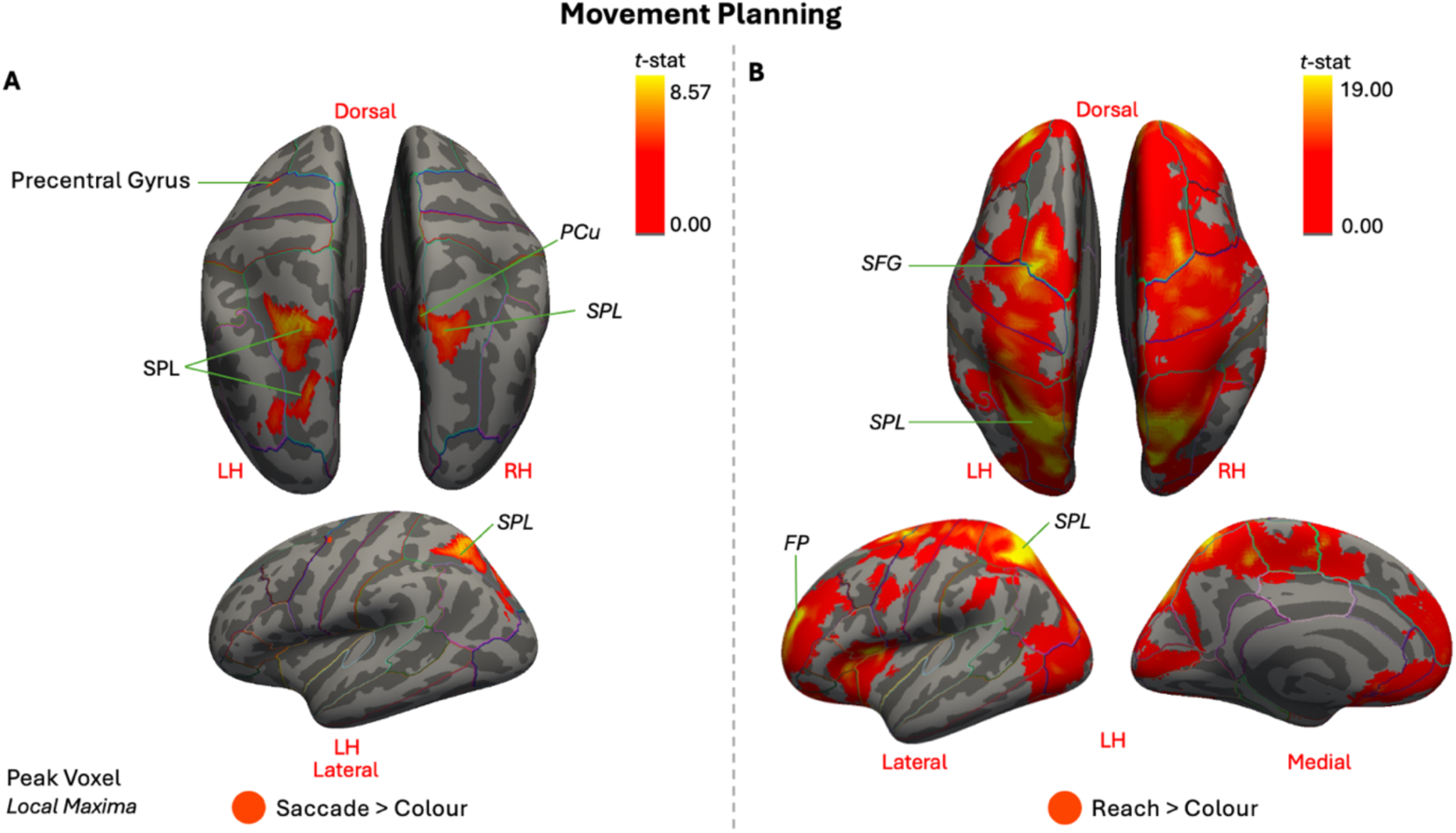
Group-level activation during the movement planning phase for the [Saccade>Colour contrast (panel A) and [Reach>Colour] contrast (panel B). Cortical results are overlaid on the FreeSurfer fsaverage ‘inflated’ brain. Light gray represents gyri and dark gray represents sulci. Lines on the surface denote the anatomical boundaries from the Desikan-Killiany Atlas, provided for visualization. Highlighted regions represent significant voxels from TFCE-corrected t-statistic maps (FWE p<0.05), with the colour-gradient corresponding to the magnitude of the t-statistic, as indicated by the colour bar. Labels are peak voxels and local maxima within clusters (local maxima shown in smaller italics).

**Table 4:** Name of region, hemisphere, MNI coordinates, t-statistic at the coordinate, and cluster sizes (number of voxels) for the movement planning phase ([Saccade>Colour] and [Reach>Colour] contrasts).

| Cluster | Region | Hemisphere | MNI<br>X | MNI<br>Y | MNI<br>Z | t-stat | Voxels |
| --- | --- | --- | --- | --- | --- | --- | --- |
| **Saccade>Colour** |  |  |  |  |  |  |  |
| 1 | SPL | Left | -18 | -62 | 56 | 8.57 | 1041 |
|  | SPL | Left | -24 | -58 | 54 | 8.08 |  |
| 2 | SPL | Right | 16 | -66 | 56 | 6.72 | 262 |
|  | PCu | Right | 8 | -60 | 58 | 6.65 |  |
| 3 | Precentral<br>Gyrus | Left | -36 | -10 | 44 | 6.03 | 22 |
| **Reach>Colour** |  |  |  |  |  |  |  |
| 1 | SPL | Left | -22 | -60 | 58 | 19 | 80445 |
|  | FP | Left | -26 | 42 | 16 | 14.4 |  |
|  | SFG | Left | -22 | -6 | 60 | 11.6 |  |

Overall, while both effectors revealed cortical engagement during this effector-dependent phase, activation differed. Saccade planning activity was relatively more focal along posterior parietal areas whereas reach planning was characterized by substantially broader and stronger activation along the cortex, with peaks in left posterior parietal and frontal regions.

#### Movement Execution

During the *movement execution* phase, participants performed the instructed movement (saccade or reach) towards the remembered target position. Consequently, the contrasts [Saccade>Colour] and [Reach>Colour] isolate regions with greater activity during effector-dependent movement, relative to the colour-report condition. **Figure 5** displays the cortical functional maps of these contrasts, overlaid on the fsaverage inflated cortical surface and **Table 5** reports the MNI coordinates and t-statistic values of peak voxels and local maxima with each significant cluster. Saccade execution (**Figure 5A**) revealed distributed occipital, posterior parietal, and frontal activation. The largest cluster’s (43982 voxels) peak activity was centered in the Occipital Fus. and extended into bilateral SPL and left PCu. Additionally, a second large cluster (10753 voxels) was centered in the left precentral gyrus, with a local maximum in right precentral gyrus. Finally, a smaller cluster revealed peak activity in the FP. In contrast, reach execution was dominated by a large cluster (69814 voxels) with activation that extended into occipital, parietal, and frontal regions, including local peak activity in the left postcentral gyrus (**Figure 5B**). Activation in this cluster was stronger in the left hemisphere. An additional smaller cluster (195 voxels) was observed more laterally, centered in the left precentral gyrus.

**Figure 5:**
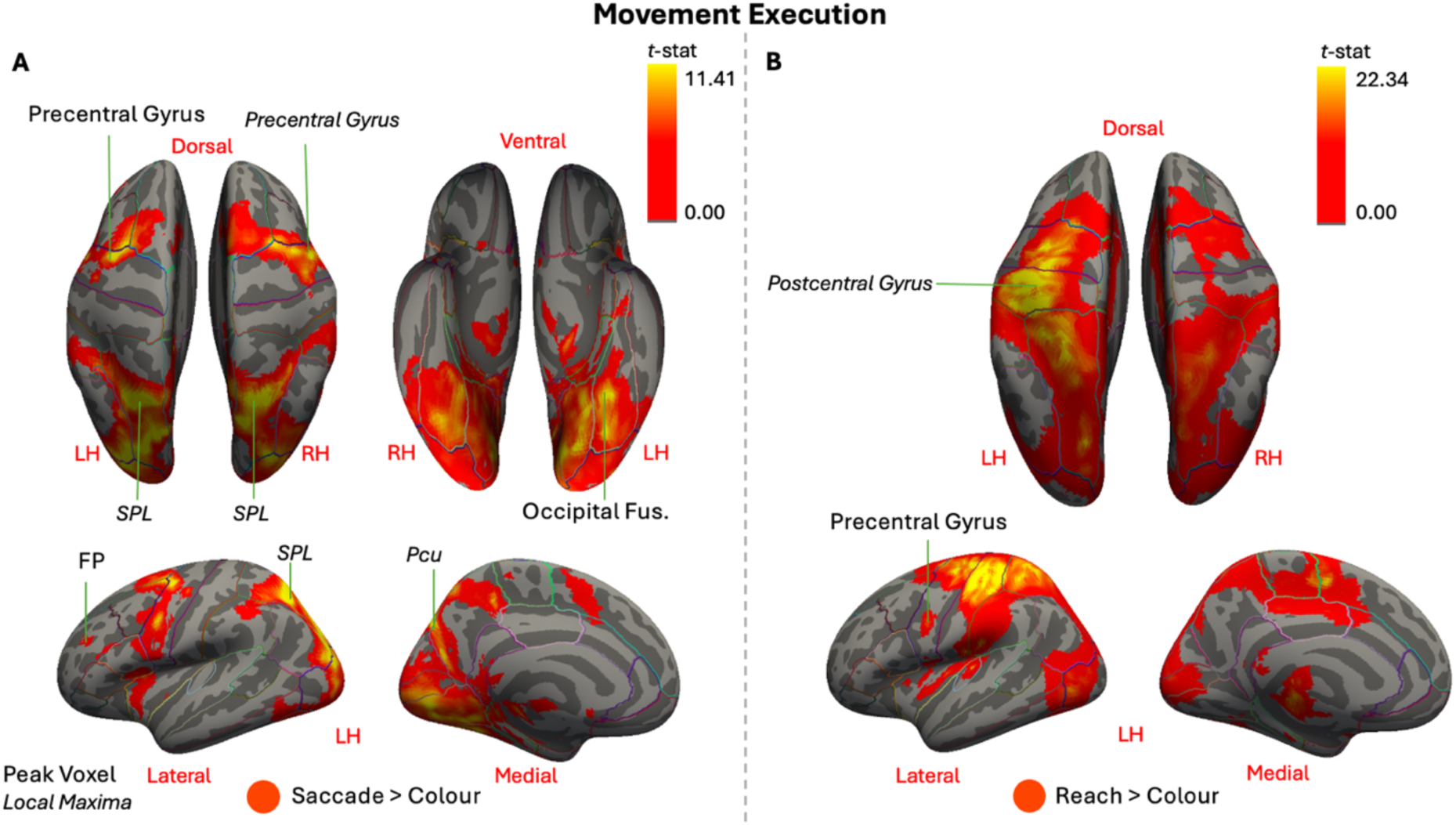
Group-level activation during the movement execution phase for the [Saccade>Colour] contrast (panel A) and [Reach>Colour] contrast (panel B). Cortical results are overlaid on the FreeSurfer fsaverage ‘inflated’ cortical surface. Light gray represents gyri and dark gray represents sulci. Lines on the surface denote the anatomical boundaries from the Desikan-Killiany Atlas, provided for visualization. Highlighted regions represent significant voxels from TFCE-corrected t-statistic maps (FWE p<0.05), with the colour-gradient corresponding to the magnitude of the t-statistic, as indicated by the colour bar. Labels are peak voxels and local maxima within clusters (local maxima shown in smaller italics).

**Table 5:**
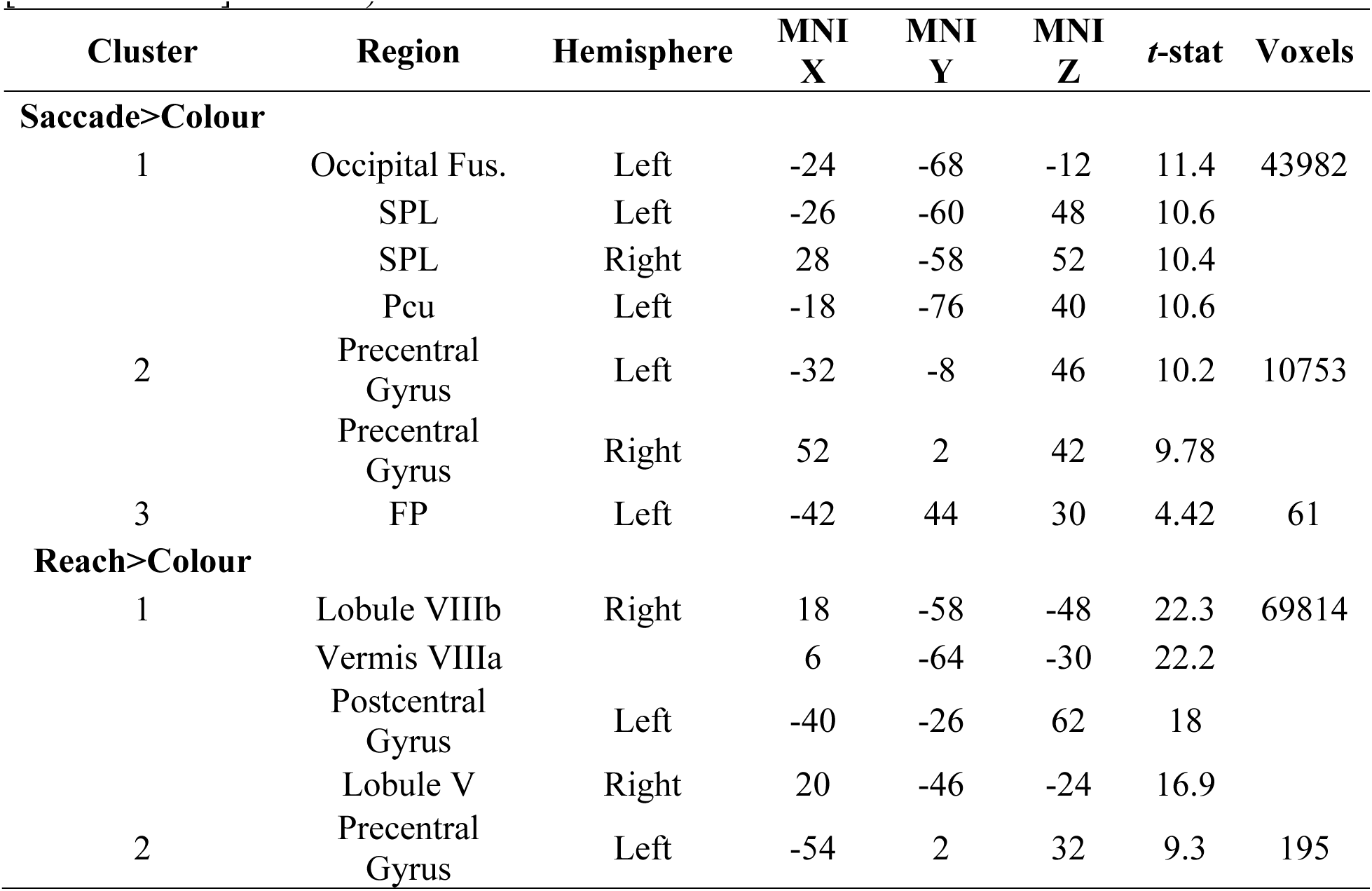
Name of region, hemisphere, MNI coordinates, t-statistic at the coordinate, and cluster sizes (number of voxels) for the movement execution phase ([Saccade>Colour] and [Reach>Colour] contrasts).

Importantly, *movement execution* phase contrasts [Saccade>Colour] and [Reach>Colour] revealed cerebellar activity, shown in **Figure 6**. During saccade execution (**Figure 6A**) there was modest cerebellar activation although no peaks were observed. In contrast, the largest cluster during reach execution (69814 voxels) had its peak activity in the right lobule V and VIIIb, as well as a local maxima extending into the vermis VIIIa (**Figure 6B**).

**Figure 6:**
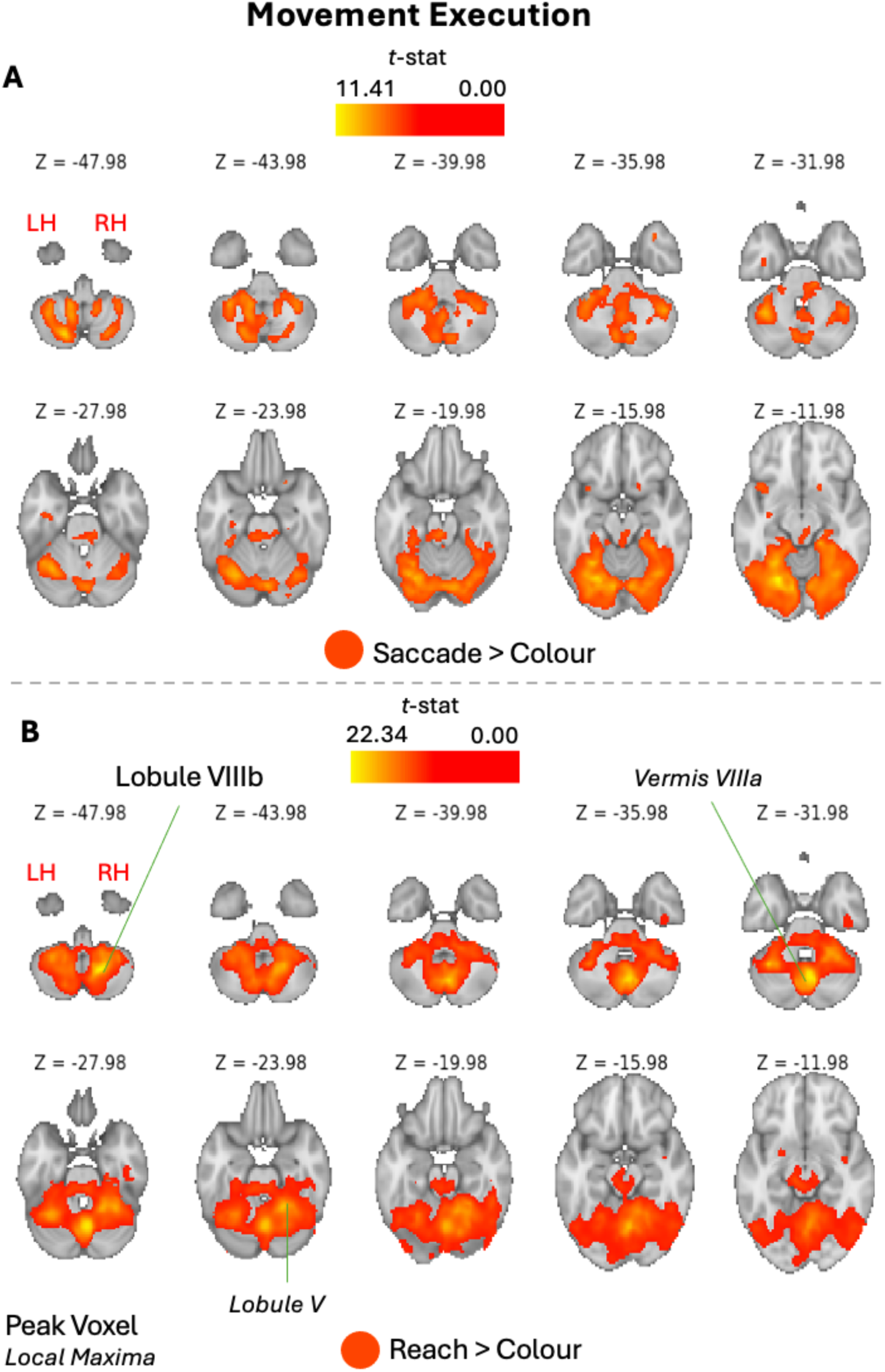
Group-level activation during the movement execution phase for the [Saccade>Colour] contrast (panel A) and [Reach>Colour] contrast (panel B). Cerebellar results are displayed on the MNI152 standard brain, shown as horizontal slices at every fourth axial z-slice in volume space. Highlighted regions represent significant voxels from TFCE-corrected t-statistic maps (FWE p<0.05), with the colour-gradient corresponding to the magnitude of the t-statistic, as indicated by the colour bar. Labels are peak voxels and local maxima within clusters (local maxima shown in smaller italics).

Altogether, both effectors produced widespread cortical activity with clear differences during the *movement execution* phase. Saccade execution was characterized with peaks in occipital, posterior parietal, and frontal areas. Although cerebellar activity was present, no peak voxels were observed. Similar to the *movement planning* phase, reach execution produced broader and stronger activity that was predominantly left-lateralized, especially along the postcentral gyrus and adjacent precentral gyrus, consistent with limb-movement related areas (Beurze et al., 2007). Reach execution also revealed peaks of the right cerebellum and the vermis.

### Overlapping saccade and reach activation during planning and execution

Our next goal was to better understand the commonalities between saccade and reach activation during the planning and execution phases.

#### Movement Planning

The *movement planning* phase for the [Saccade>Colour] contrast revealed peak activity along bilateral posterior parietal regions and in the left precentral regions, whereas the [Reach>Colour] contrast showed more widespread activity, with peaks posterior parietal and frontal regions (**Figure 5**). The overlap of these two maps ([Saccade>Colour] ∩ [Reach>Colour]) is presented in **Figure 7**, overlaid on the fsaverage inflated cortical surface. The most immediate observation is the strong similarity between the overlapping and [Saccade>Colour] contrast maps. Notably, all significant peak voxels and local maxima identified during saccade planning were present in this conjunction map. Thus, overlapping activation during reach and saccade planning is observed bilaterally in SPL, as well as in the right PCu and left precentral gyrus. Importantly, the results here simply identify voxels that were significant in both contrasts, but it does not provide information about the relative strength of activation.

**Figure 7:**
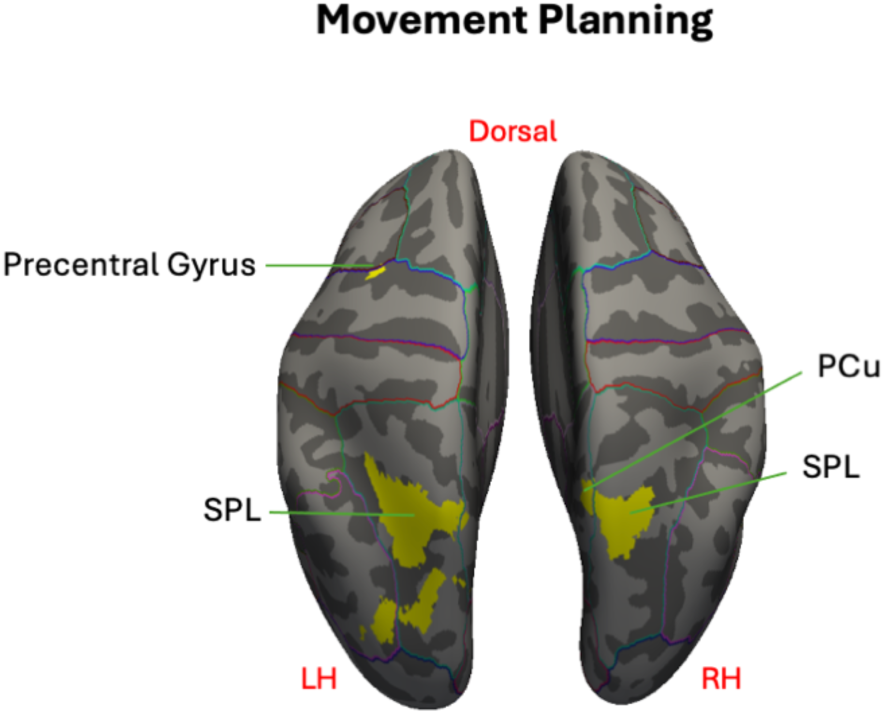
Overlapping activation for reach and saccade during the movement planning phase ([Saccade>Colour] ∩ [Reach>Colour]). Cortical results are overlaid on the FreeSurfer fsaverage ‘inflated’ cortical surface. Light gray represents gyri and dark gray represents sulci. Lines on the surface denote the anatomical boundaries from the Desikan-Killiany Atlas, provided for visualization. Highlighted regions represent voxels significant in both contrasts, suggesting possible involvement for both effectors. Coordinates for labels come from peak voxels and local maxima of [Saccade>Colour] contrast during the movement planning phase.

In summary, overlapping activation for the saccade and reach condition was mostly revealed along bilateral posterior parietal areas during the *movement planning* phase. The spatial pattern of overlap closely matched what was observed during saccade planning while the broader cortical activity present during reach planning was not retained.

#### Movement Execution

During movement execution, the [Saccade>Colour] contrast revealed activity along occipital, posterior parietal, and frontal regions (**Figure 5A**). The [Reach>Colour] contrast revealed widespread activation along cerebellar and cortical areas, with peak activity localized within the posterior cerebellum, left postcentral gyrus and precentral gyrus (**Figure 5B**; **Figure 6B**). The cortical overlap of these two maps ([Saccade>Colour] ∩ [Reach>Colour]) is presented in **Figure 8**, overlaid on the fsaverage inflated cortical surface. Similar to the *movement planning* phase, overlapping cortical activation was similar to the functional map produced for saccade execution, with all significant peak voxels and local maxima identified during saccade execution present in this conjunction map. Thus, saccade and reach execution both engage the bilateral SPL, precentral gyrus, as well the left PCu.

**Figure 8:**
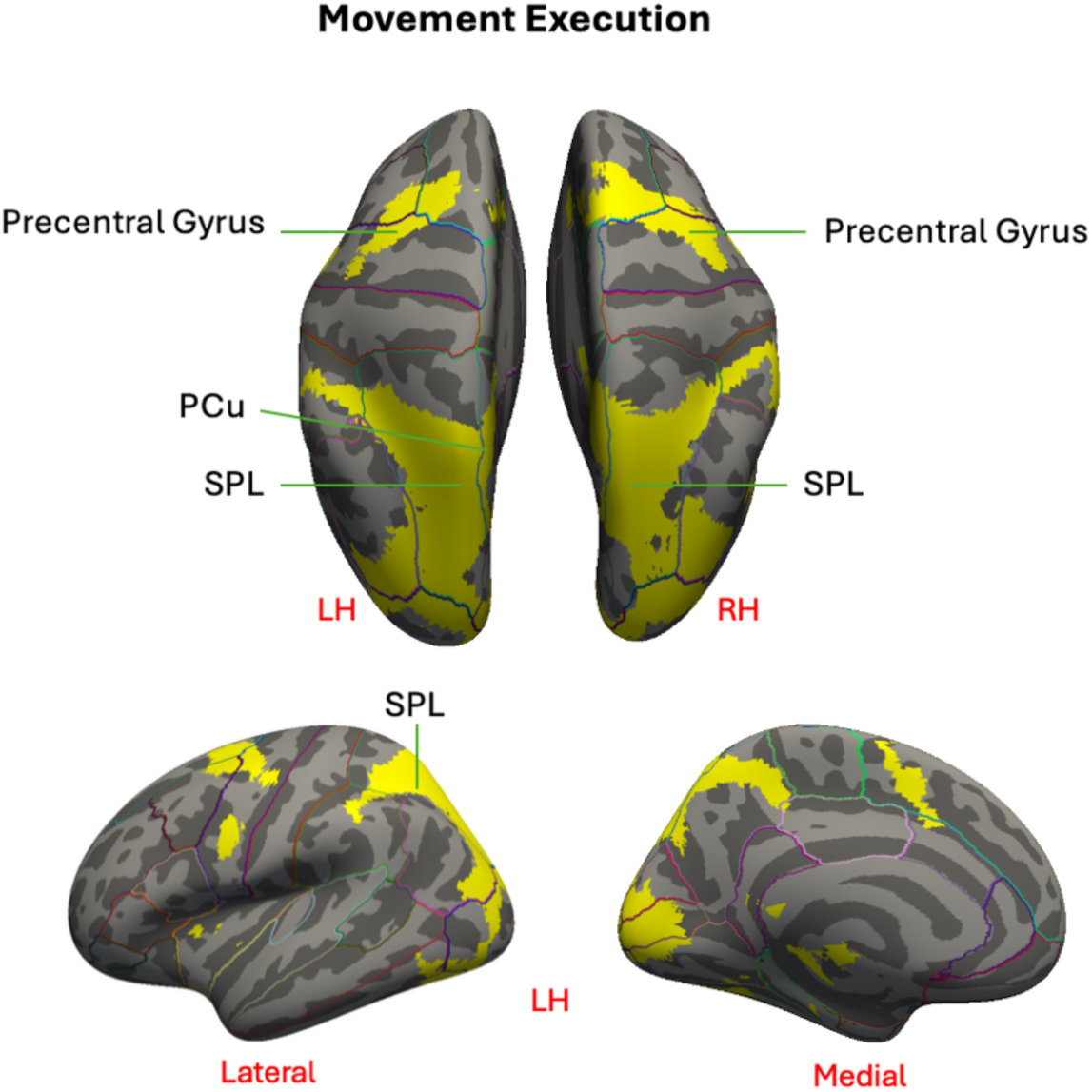
Overlapping activation for reach and saccade during the movement execution phase ([Saccade>Colour] ∩ [Reach>Colour]). Cortical results are overlaid on the FreeSurfer fsaverage ‘inflated’ cortical surface. Light gray represents gyri and dark gray represents sulci. Lines on the surface denote the anatomical boundaries from the Desikan-Killiany Atlas, provided for visualization. Highlighted regions represent voxels significant in both contrasts, suggesting possible involvement for both effectors. Coordinates for labels come from peak voxels and local maxima of [Saccade>Colour] contrast during the movement execution phase.

There was also overlap within cerebellar regions, shown in **Figure 9**, although engagement did not coincide with any of the peak and local maxima coordinates present within the [Reach>Colour] contrast, despite some overlap within the same anatomical regions. As a reminder, although activation did expand to cerebellar regions in the [Saccade>Colour] functional map, no peak voxels or local maxima were identified. Visual inspection of the conjunction map revealed overlapping activity within bilateral cerebellar lobules, including but not limited to lobule VI, Crus I, and lobule VIIIa, extending medially into the vermal VI and VIIIa (labelled in **Figure 9** for visualization). Once again, this analysis only provides information about areas that were significantly active in both contrasts but does not account for the strength of activation during saccade and reach execution.

**Figure 9:**
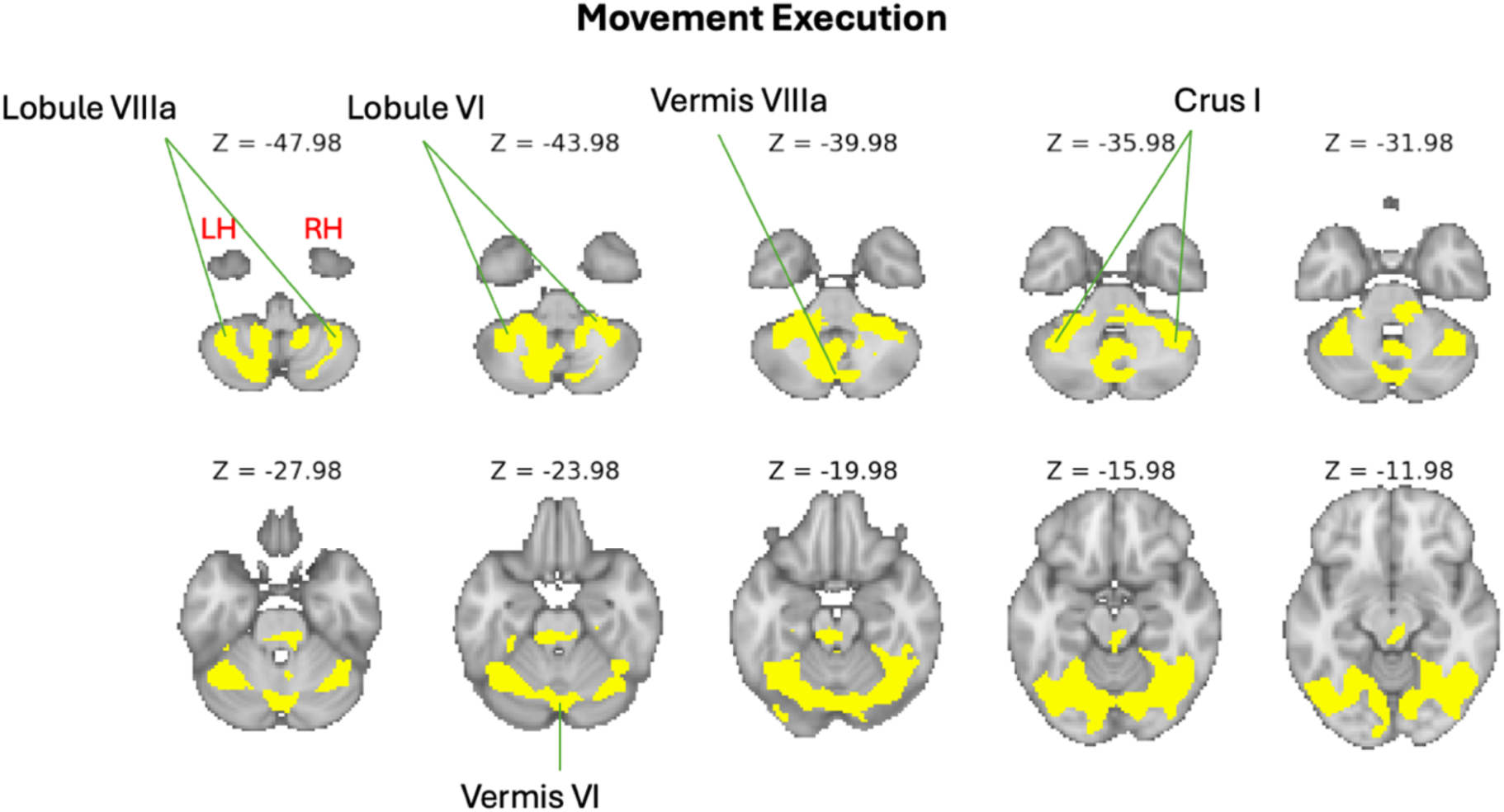
Overlapping activation for reach and saccade during the movement execution phase ([Look>Colour] ∩ [Reach>Colour]). Cerebellar results are displayed on the MNII152 standard brain, shown as horizontal slices at every fourth axial z-slice in volume space. Highlighted regions represent significant voxels from TFCE-corrected t-statistic maps (FWE p<0.05), with the colour-gradient corresponding to the magnitude of the t-statistic, as indicated by the colour bar. Labels are provided for visualization but do not correspond to peak voxels or local maxima of [Saccade>Colour] or [Reach>Colour] contrasts.

Overall, the *movement execution* phase showed overlapping activity between saccade and reach conditions in bilateral occipital, posterior parietal, and frontal areas, as well as broad cerebellar regions. Similar to *movement planning*, the overlap map closely resembled what was observed for saccade execution, although the more extensive activation observed during reach execution was not retained.

### Effector-preferential activation across planning and execution

To determine regions with preferential activation for reach versus saccade movements, whole-brain contrasts that directly compared effectors were performed for the effector-dependent planning (contrasts 4 and 5) and execution phases (contrasts 8 and 9).

#### Movement planning

Direct [Saccade>Reach] and [Reach>Saccade] contrasts during the *movement planning* phase identified regions that are more strongly engaged for saccades versus reaches and vice versa, respectively. Functional maps of these contrasts, overlaid on the fsaverage inflated cortical surface, are presented in **Figure 10**. **Table 6** reports the MNI coordinates and t-statistic values of peak voxels and local maxima with each significant cluster. Saccade-preferential activation was characterized by a large cluster (15021 voxels) centered in the ventral occipitotemporal cortex. The peak voxel was centered in the left Temporal Fus. (**Figure 10A**), with activation extending into the right Temporal Fus. An additional, smaller cluster (208 voxels) was centered in the right LOC. In contrast, reach-preferential activation was dominated by a single, very large cluster (83599 voxels) that spanned bilateral occipital, parietal, and frontal regions. Peak activity for this cluster was present within the left SPL and extended to a local maximum in the left LOC (**Figure 16B**).

**Figure 10:**
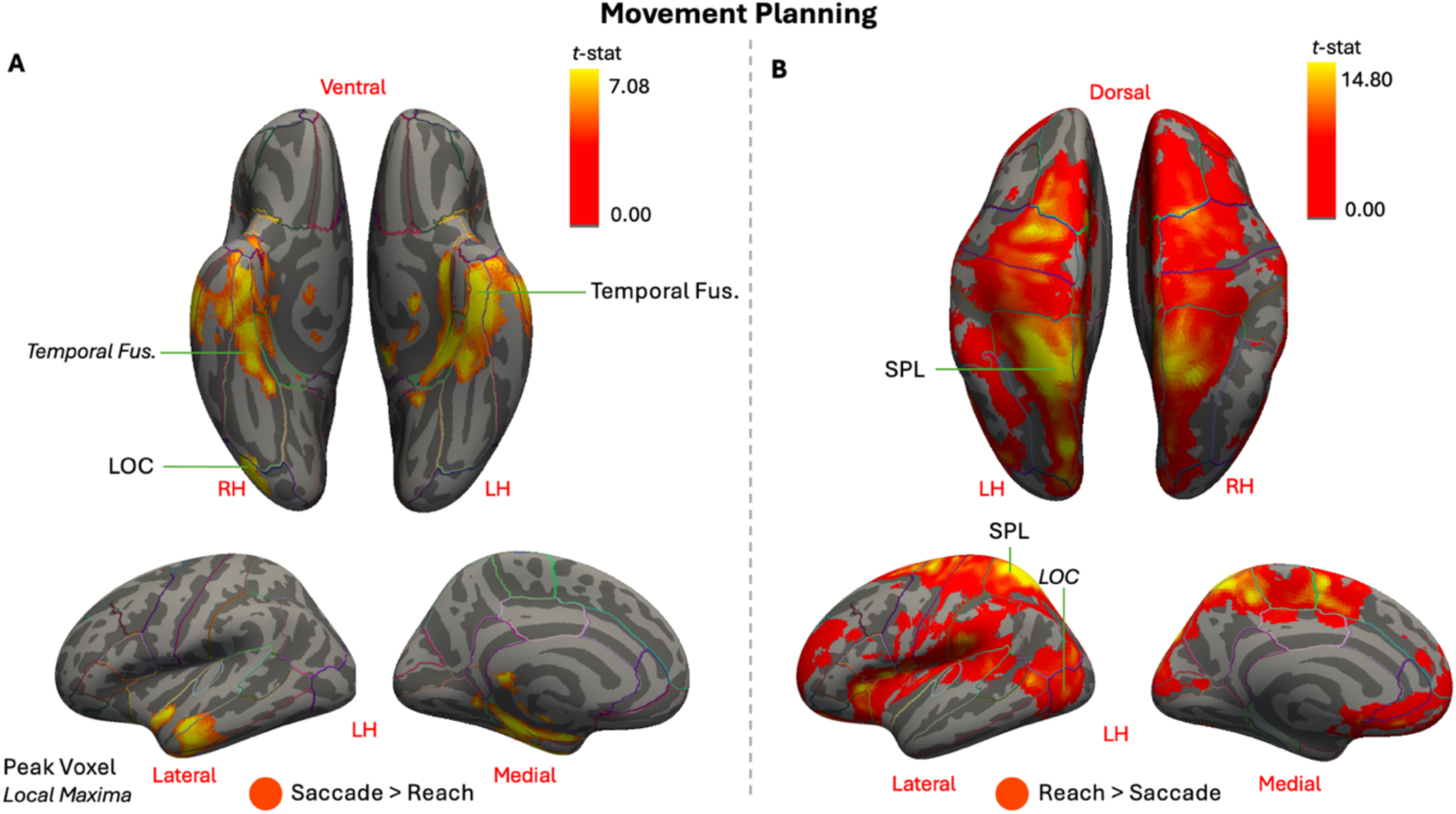
Group-level activation during the movement planning phase for the [Saccade>Reach] contrast (panel A) and [Reach>Saccade] contrast (panel B). Cortical results are overlaid on the FreeSurfer fsaverage ‘inflated’ cortical surface. Light gray represents gyri and dark gray represents sulci. Lines on the surface denote the anatomical boundaries from the Desikan-Killiany Atlas, provided for visualization. Highlighted regions represent significant voxels from TFCE-corrected t-statistic maps (FWE p<0.05), with the colour-gradient corresponding to the magnitude of the t-statistic, as indicated by the colour bar. Labels are peak voxels and local maxima within clusters (local maxima shown in smaller italics).

**Table 6:** Name of region, hemisphere, MNI coordinates, t-statistic at the coordinate, and cluster sizes (number of voxels) for the movement planning phase ([Saccade>Reach] and [Reach>Saccade] contrasts).

| Cluster | Region | Hemisphere | MNI<br>X | MNI<br>Y | MNI<br>Z | t-stat | Voxels |
| --- | --- | --- | --- | --- | --- | --- | --- |
| **Saccade>Reach** |  |  |  |  |  |  |  |
| 1 | Temporal Fus. | Left | -38 | -6 | -28 | 7.08 | 15021 |
|  | Temporal Fus. | Right | 30 | -32 | -24 | 6.18 |  |
|  | Lobule I-IV | Left | -6 | -42 | -18 | 6.16 |  |
| 2 | LOC | Right | 42 | -76 | -12 | 6.32 | 208 |
| **Reach>Saccade** |  |  |  |  |  |  |  |
| 1 | SPL | Left | -22 | -60 | 60 | 14.8 | 83599 |
|  | LOC | Left | -16 | -84 | 40 | 10.1 |  |

Cerebellar activation for saccade-preferential and reach-preferential areas is shown in **Figure 11**. Saccade-related activity from cluster 1 extended into the left lobule I-IV (**Figure 11A**) while no reach-preferential cerebellum activity was observed during this phase (**Figure 11B**).

**Figure 11:**
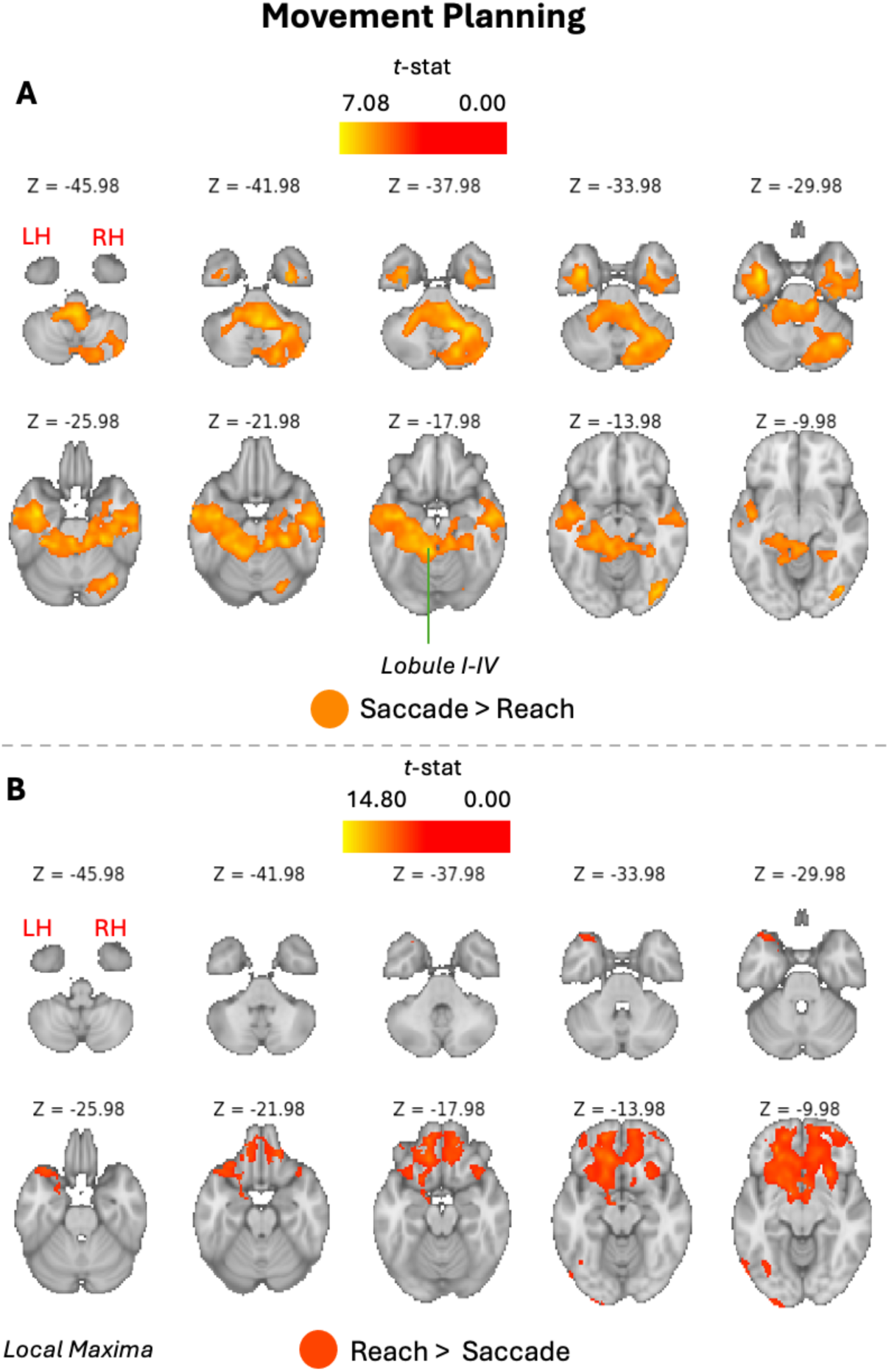
Group-level activation during the movement planning phase for the [Saccade>Reach] contrast (panel A) and [Reach>Saccade] contrast (panel B). Subcortical results are displayed on the MNII152 standard brain, shown as horizontal slices at every fourth axial z-slice in volume space. Highlighted regions represent significant voxels from TFCE-corrected t-statistic maps (FWE p<0.05), with the colour-gradient corresponding to the magnitude of the t-statistic, as indicated by the colour bar. Labels are peak voxels and local maxima within clusters (local maxima shown in smaller italics).

Overall, saccade– and reach-preferential activity was revealed during the planning phase, with distinct spatial organizations. Saccade-preferential activity was evident in the ventral occipitotemporal cortex as well as the left cerebellum while reach-preferential activity produced broader and stronger cortical activity spanning dorsal occipital and posterior parietal regions.

#### Movement Execution

Similar to the *movement planning* phase, direct contrasts during movement execution identified regions with greater activity for saccades versus reaches, and vice versa. Functional maps for the [Saccade>Reach] and [Reach>Saccade] contrasts, overlaid on the fsaverage inflated cortical surface, are presented in **Figure 12**. **Table 7** reports the MNI coordinates and t-statistic values of peak voxels and local maxima with each significant cluster. Saccade-preferential activation was dominated by a large cluster (78018 voxels) with an activation pattern that spanned occipital, lateral parietal, and superior and middle frontal regions (**Figure 12A**). The peak voxel was located in the left Occipital Fus, with activation extending into left AG/SMG, left LOC, and left LG. Conversely, reach-preferential activation was present in a large cluster (38205 voxels) exhibiting a spatial pattern that spanned occipital, medial parietal, and frontal regions (**Figure 12B**). Local peak activity was found in the left postcentral gyrus. A smaller cluster (543 voxels) was also present, centered in the left LOC.

**Figure 12:**
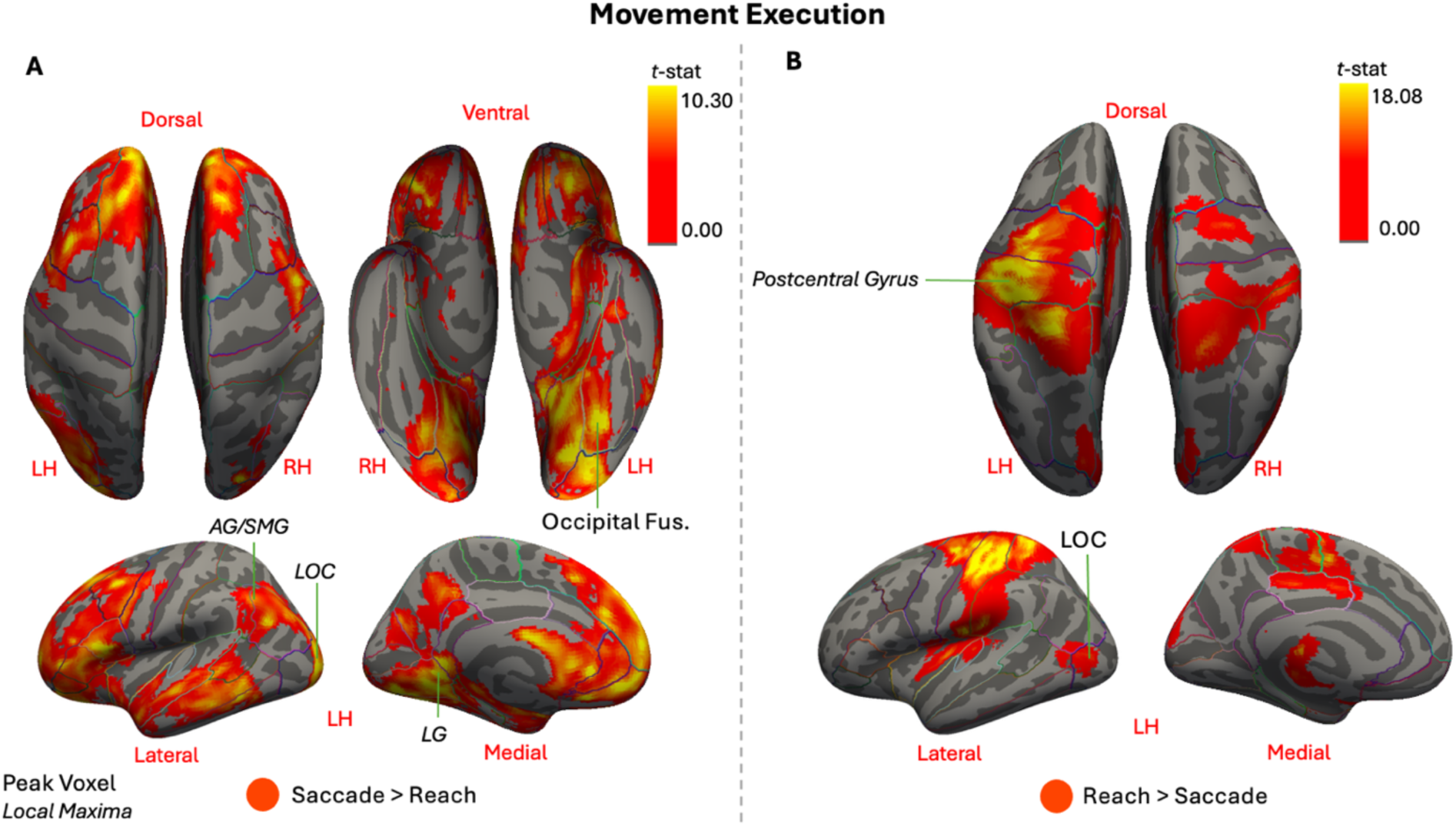
Group-level activation during the movement execution phase for the [Saccade>Reach] contrast (panel A) and [Reach>Saccade]) contrasts (panel B). Cortical results are overlaid on the FreeSurfer fsaverage ‘inflated’ cortical surface. Light gray represents gyri and dark gray represents sulci. Lines on the surface denote the anatomical boundaries from the Desikan-Killiany Atlas, provided for visualization. Highlighted regions represent significant voxels from TFCE-corrected t-statistic maps (FWE p<0.05), with the colour-gradient corresponding to the magnitude of the t-statistic, as indicated by the colour bar. Labels are peak voxels and local maxima within clusters (local maxima shown in smaller italics).

**Table 7:**
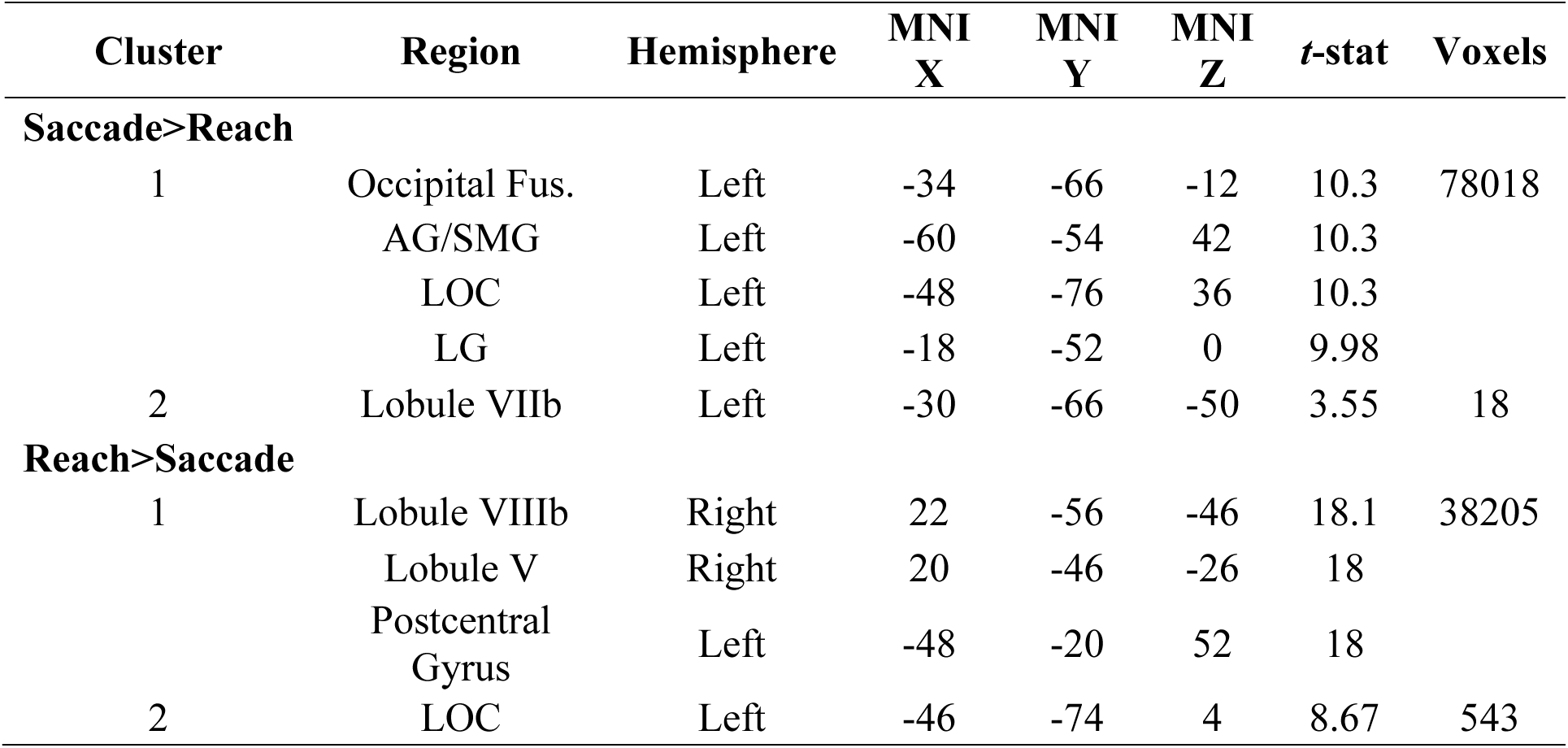
Name of region, hemisphere, MNI coordinates, t-statistic at the coordinate, and cluster sizes (number of voxels) for the movement execution phase ([Saccade>Reach] and [Reach>Saccade] contrasts).

| Cluster | Region | Hemisphere | MNI<br>X | MNI<br>Y | MNI<br>Z | t-stat | Voxels |
| --- | --- | --- | --- | --- | --- | --- | --- |
| **Saccade>Reach** |  |  |  |  |  |  |  |
| 1 | Occipital Fus. | Left | -34 | -66 | -12 | 10.3 | 78018 |
|  | AG/SMG | Left | -60 | -54 | 42 | 10.3 |  |
|  | LOC | Left | -48 | -76 | 36 | 10.3 |  |
|  | LG | Left | -18 | -52 | 0 | 9.98 |  |
| 2 | Lobule VIIb | Left | -30 | -66 | -50 | 3.55 | 18 |
| **Reach>Saccade** |  |  |  |  |  |  |  |
| 1 | Lobule VIIIb | Right | 22 | -56 | -46 | 18.1 | 38205 |
|  | Lobule V | Right | 20 | -46 | -26 | 18 |  |
|  | Postcentral Gyrus | Left | -48 | -20 | 52 | 18 |  |
| 2 | LOC | Left | -46 | -74 | 4 | 8.67 | 543 |

**Figure 13** shows the cerebellar activation. Saccade-preferential activation in cluster 1 expanded into the cerebellum, although peak activity was not found here (**Figure 13A**). A smaller, focal cluster (18 voxels) was also present in the left lobule VIIb (not labelled). Conversely, reach-preferential activation in the largest revealed peak activity in the right lobule VIIIb, with a local maximum in the right lobule V (**Figure 13B**).

**Figure 13:**
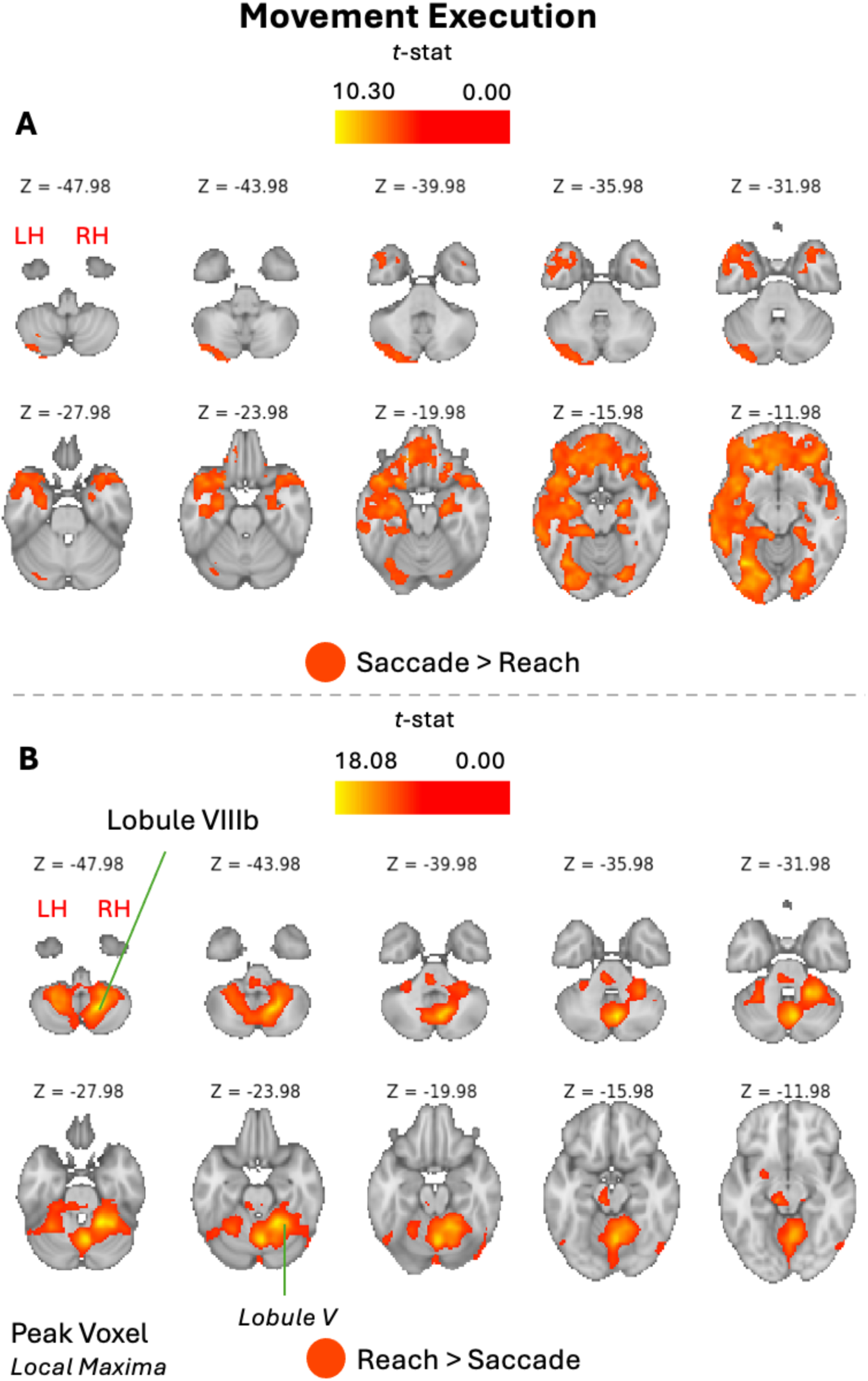
Group-level activation during the movement execution phase for the [Saccade>Reach] contrast (panel A) and [Reach>Saccade]) contrast (panel B). Subcortical results are displayed on the MNI152 standard brain, shown as horizontal slices at every fourth axial z-slice in volume space. Highlighted regions represent significant voxels from TFCE-corrected t-statistic maps (FWE p<0.05), with the colour-gradient corresponding to the magnitude of the t-statistic, as indicated by the colour bar. Labels are peak voxels and local maxima within clusters (local maxima shown in smaller italics).

In summary, effector-preferential differences were most pronounced during the *movement execution* phase. Saccade-preferential activity was concentrated mostly along ventral occipital and lateral posterior parietal regions, while reach-preferential activity showed a more medial and dorsal distribution spanning parietal and frontal spatial cortices, with peak activity in the left postcentral gyrus and right cerebellum. These patterns align with those observed during saccade and reach execution, respectively (**Figures 5-6**), where reach execution similarly revealed peak activity in the left postcentral gyrus and right cerebellum while saccade execution did not.

However, through directly contrasting reach and saccade conditions, more medial-lateral spatial patterns emerged for activation along the PPC.

### Region-of-interest analysis

The comparison of saccade and reach activity with the colour-report condition revealed several occipital, parietal, and frontal regions exhibiting effector-dependent activity during planning and execution phases. Additionally, direct contrasts between these effectors identified cortical regions showing greater activation for reach versus saccade, and vice versa. However, to what extent do cortical regions show differences in activation between saccade and reach? To quantify these differences, a ROI analysis was conducted by comparing BOLD percent signal change for the [Saccade>Colour] and [Reach>Colour] contrasts within functionally defined regions (**Table 1**) consistently implicated in the literature for reach and saccade processing.

#### Movement Planning

Data shown in **Figure 14** illustrate the ROI analysis for the *movement planning* phase. Significant difference (paired *t*-test; FDR-corrected p<0.05) between saccade– and reach-related activity was observed in the right aCing, bilateral DLPFC, MedFG, bilateral MidFG, bilateral PMd, bilateral M1, left AG, left mIPS, and bilateral SPOC (**Figure 14A**). Of these, activity was significantly greater for saccades than the colour condition only in left mIPS, and significantly lower in left M1, left MidFG, MedFG, and left S1 (one-sample *t*-test, FDR-corrected p<0.05; **Figure 14B**). Reach activity was significantly greater than colour in all regions shown in **Figure 14B**, except left AG, right SPOC, left MidFG, and left S1. Critically, all of these areas showed significantly greater activation values for reach than saccade, indicating reach-preferential activity within these areas during movement preparation. This preference comes from the higher relative mean BOLD signal change for reaches compared with saccades, despite some regions showing larger effect magnitudes for saccades.

**Figure 14:**
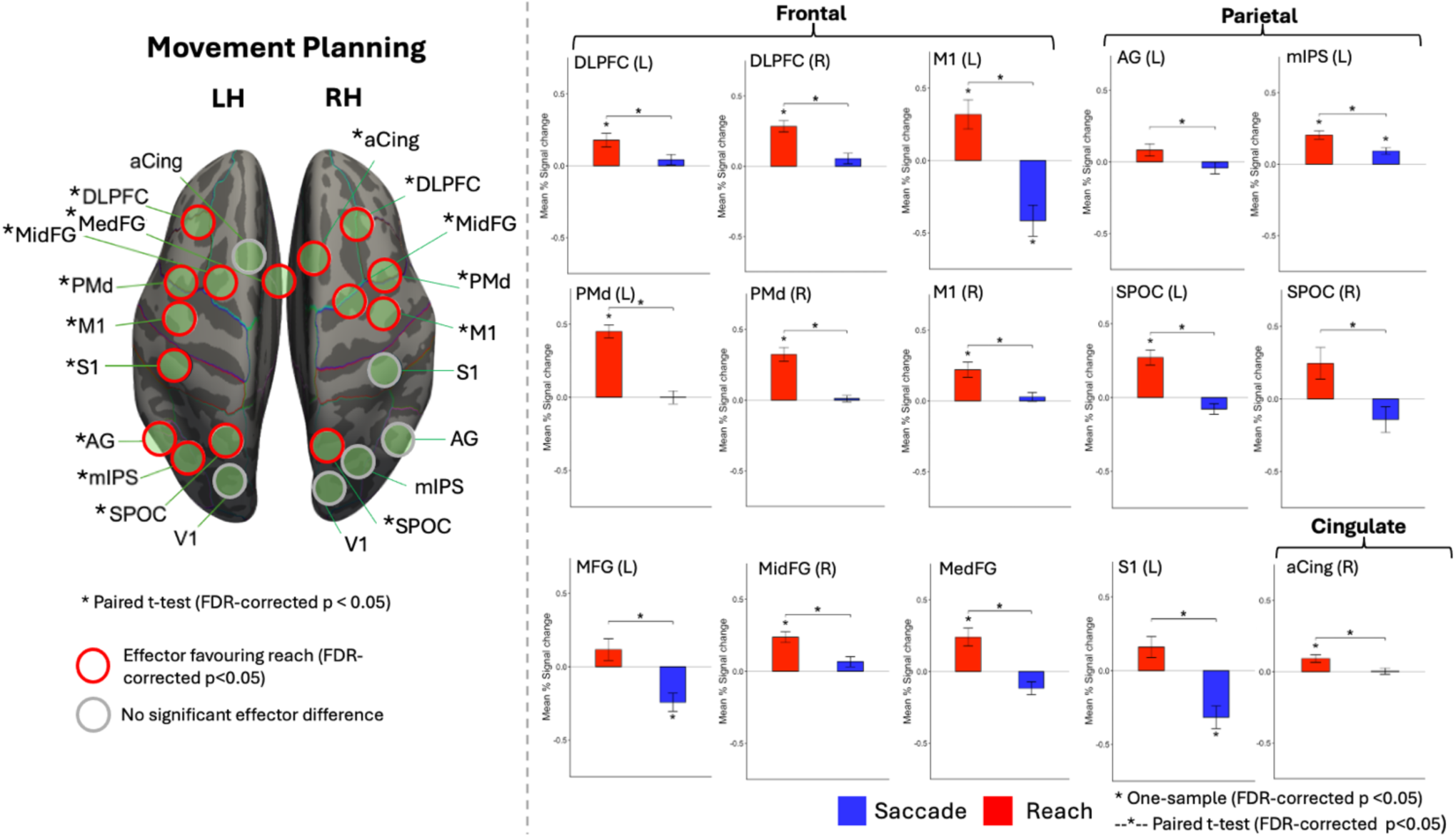
ROI analysis for selected regions during the movement planning phase. ROIs are displayed on the FreeSurfer fsaverage ‘inflated’ cortical surface (panel A). Asterisks beside ROI labels indicate a significant difference between the [Saccade>Colour] and [Reach>Colour] contrasts (paired *t*-test; FDR-corrected p<0.05). Red outlines identify ROIs with significantly greater mean % signal change values for reach compared to saccade, whereas gray outlines represent no significant difference between effectors. Mean % signal change for ROIs showing a significant effector difference is plotted (panel B). Asterisks above bars indicate significant one-sample effects (FDR-corrected p<0.05), and brackets denote significant differences between effectors from paired t-tests (FDR-corrected p<0.05). Red bars correspond to reach values and blue bars to saccade. Error bars represent SEM.

#### Movement Execution

**Figure 15** reveals the ROI analysis for the *movement execution* phase. Significant difference (paired *t*-test; FDR-corrected p<0.05) between saccade– and reach-related activity was observed in the right aCing, left DLPFC, MedFG, left MidFG, bilateral PMd, left M1, left AG, left mIPS, bilateral SPOC, and right V1 (**Figure 15A**). Of these, activity was significantly greater for saccades than the colour condition in left mIPS, right V1, left SPOC and left PMd, and lower in left MidFG, left M1, and left S1 (one-sample *t*-test, FDR-corrected p<0.05; **Figure 15B**). Reach activity was significantly greater than the colour condition in all areas shown in **Figure 15B** except left AG, where activity was significantly lower. In contrast to the *movement planning* phase, several regions showed significant differences favoring saccades, including left DLPFC, left AG, left mIPS, and right V1. However, note that in right DLPFC neither effector showed significant activity greater than the colour condition and in left AG both saccade– and reach-related activity were significantly lower than the colour condition, with saccades being significantly less negative than reach. All other significant regions favored reach-related activity.

**Figure 15:**
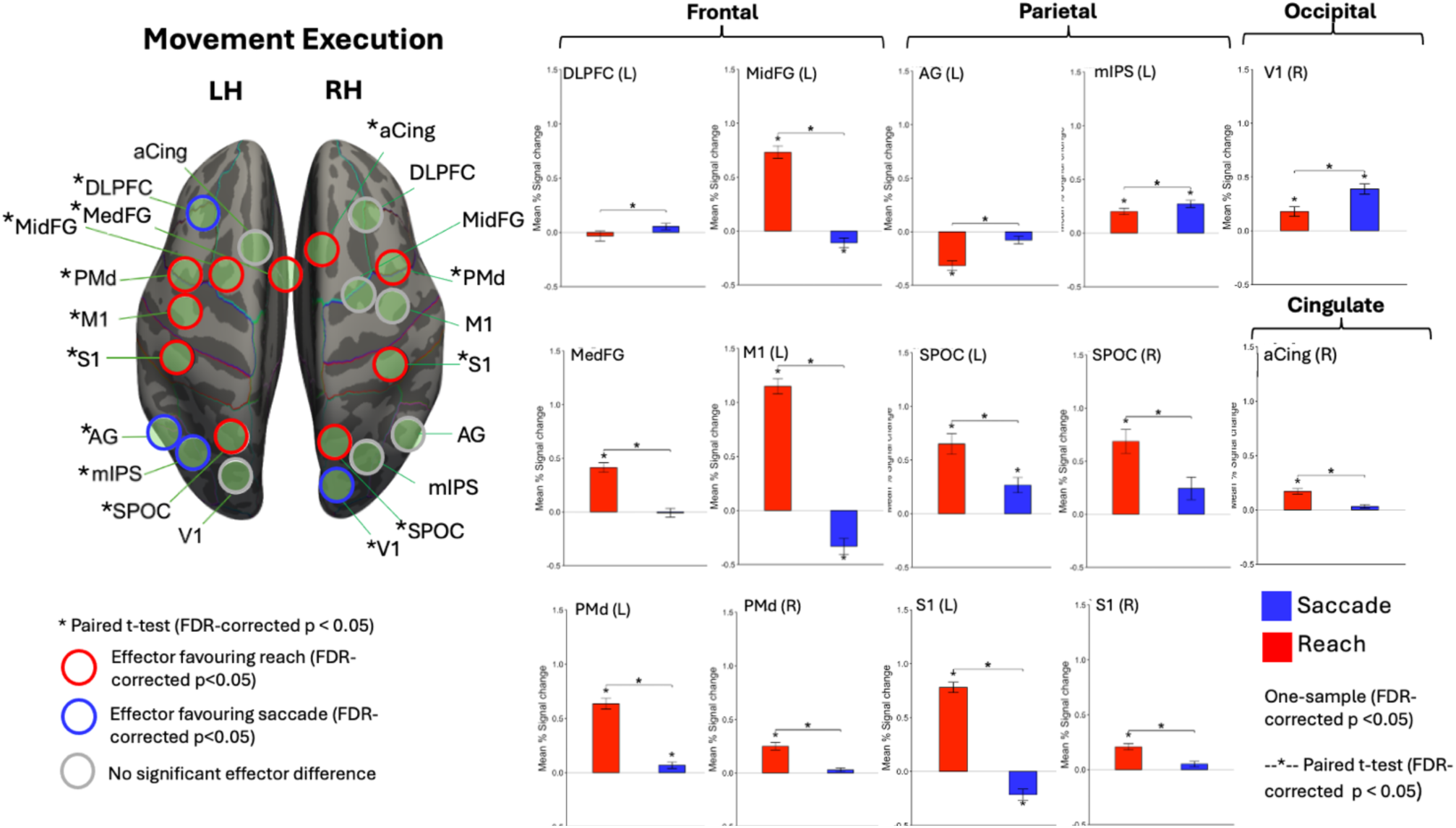
ROI analysis for selected regions during the movement execution phase. ROIs are displayed on the FreeSurfer fsaverage ‘inflated’ cortical surface (panel A). Asterisks beside ROI labels indicate a significant difference between the [Saccade>Colour] and [Reach>Colour] contrasts (paired *t*-test; FDR-corrected p<0,05). Red outlines identify ROIs with significantly greater mean % signal change values for reach compared to saccade, blue outlines identify ROIs with significantly greater mean % signal change for saccade compared to reach, and gray outlines represent no significant difference between effectors. Mean % signal change for ROIs showing a significant effector difference is plotted (panel B). Asterisks above bars indicate significant one-sample effects (FDR-corrected p<0.05), and brackets denote significant differences between effectors from paired t-tests (FDR-corrected p<0.05). Red bars correspond to reach values and blue bars to saccade. Error bars represent SEM.

### Multivariate Analysis

To determine whether saccade and reach planning and execution were associated with distinct multivoxel activity patterns, we calculated crossvalidated squared Euclidean distances between saccade and reach representations within each ROI.

During **planning**, significant saccade-reach multivoxel dissimilarity was observed in 16 of 28 ROIs using raw parameter estimates (**Figure 16, top**). These included bilateral PMd, S1, FEF, and M1, as well as right SPOC, right mIPS, SEF, right aCing, left cerebellar Crus I, right cerebellar lobule VI, left cerebellar lobule VIIIb, and vermis VI (all FDR-adjusted p ≤ .034). The strongest statistical evidence for dissimilarity was observed in right PMd (t(18) = 4.63, pFDR = .0059), followed by right FEF (t(18) = 3.64, pFDR = .0110).

Following removal of the within-ROI mean activity for each condition and run, significant saccade-reach distances were observed in only 6 of 28 ROIs during planning (**Figure 16, top**). These included bilateral FEF, right PMd, right mIPS, right aCing, and right cerebellar lobule VI (all FDR-adjusted p ≤ .035). Whereas distances calculated from raw parameter estimates may reflect differences in both overall activation magnitude and spatial activity patterns, voxel-mean removal eliminates the spatially uniform component of the response.

**Figure 16.**
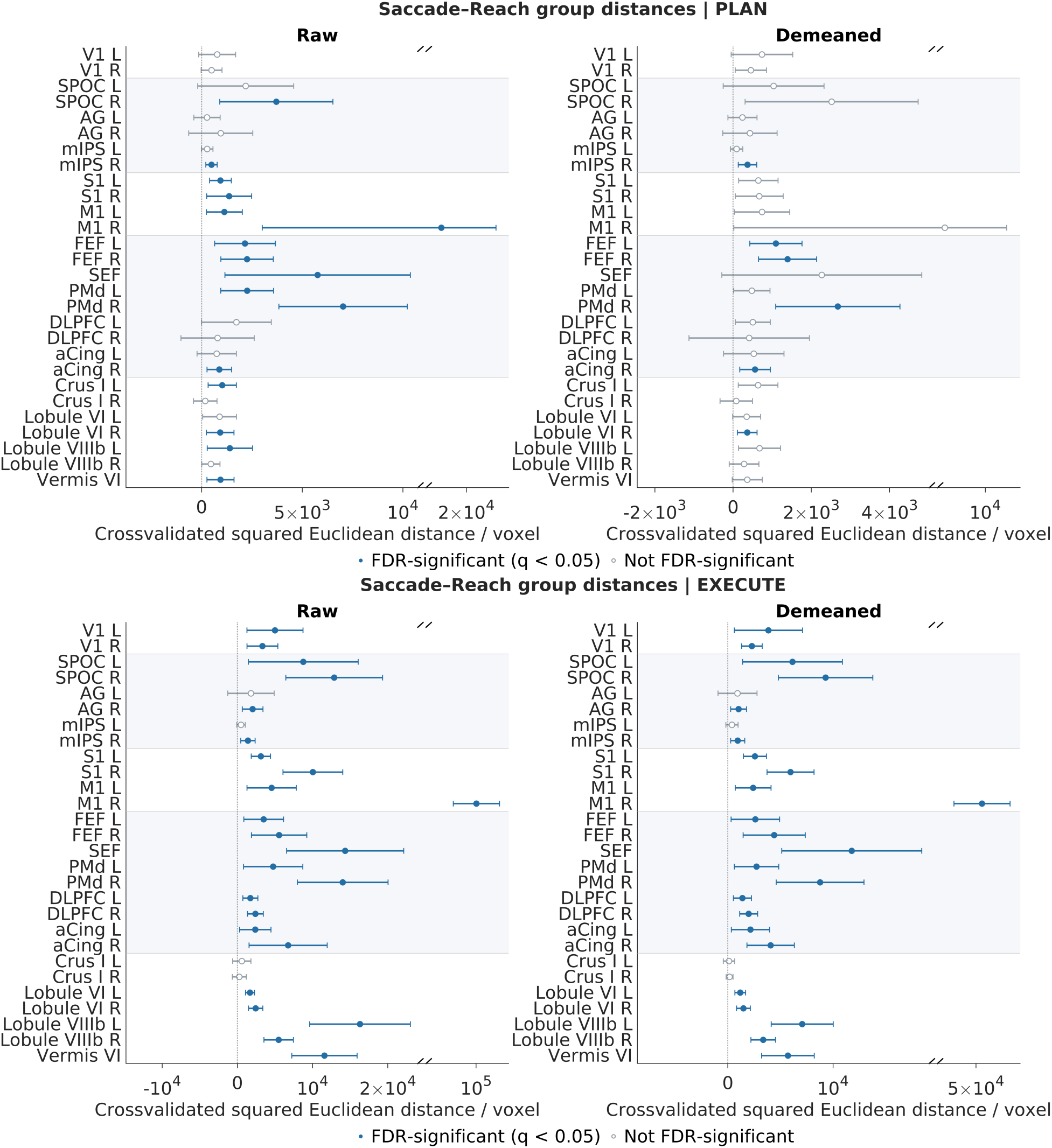
Saccade-reach representational dissimilarity across cortical and cerebellar regions. Top panel: Plan phase. Bottom panel: Execute phase. Circles represent group means, with horizontal error bars indicating 95% confidence intervals. Blue filled circles indicate ROIs with significant representational distances following FDR correction (q < 0.05), whereas grey open circles indicate nonsignificant ROIs. ROIs are grouped anatomically into occipital, parietal, sensorimotor, frontal, and cerebellar regions. The vertical dotted line indicates zero representational distance. The axis includes a scale break to compress the larger M1-R distances and improve visualization of the remaining ROIs.

During **execution**, significant saccade-reach distances were observed in the same 24 of 28 ROIs using both raw and demeaned parameter estimates (**Figure 16, bottom**), indicating that effector-dependent differences remained widespread following removal of mean regional activation. These included widespread cortical and cerebellar regions, notably bilateral PMd, M1, S1, FEF, SPOC, DLPFC, V1, and aCing, as well as bilateral cerebellar lobules VI and VIIIb and vermis VI.

## DISCUSSION

The present study employed a cue-separated event-related fMRI paradigm to investigate four questions: (1) how saccade– and reach-related activity evolves across visual target encoding, movement planning, and execution; (2) which cortical and cerebellar regions exhibit overlapping activation between effectors; (3) which regions demonstrate effector-preferential activity; and (4) whether saccades and reaches elicit distinguishable multivoxel activity patterns within these regions.

First, visual target encoding primarily recruited frontal and temporal regions, whereas effector-dependent activity emerged during movement planning, with saccades recruiting posterior parietal regions and reaches engaging a broader occipito-parieto-frontal network. During execution, both cortical and cerebellar networks were extensively engaged. Second, overlapping activation was observed across parieto-frontal regions during planning and execution, with additional cerebellar overlap. Third, direct contrasts revealed limited saccade-preferential activity in ventral occipital and cerebellar regions and more widespread reach-preferential activity during planning. During execution, a relative medial-lateral organization of reach– and saccade-preferential activity emerged, respectively, alongside reach-preferential activation in the ipsilateral cerebellum (right lobules V and VIIIb). ROI analyses further identified predominantly greater activation for reaches, particularly across frontal and parietal regions.

Finally, multivariate analyses extended these findings by demonstrating reliably distinguishable saccade– and reach-related activity patterns across widespread cortical and cerebellar regions during both planning and execution, including after removing differences in mean regional activation. Together, these findings indicate that although saccades and reaches recruit overlapping cortical and cerebellar regions, their spatial activity patterns can remain distinguishable, highlighting the complementary contributions of univariate and multivariate analyses to understanding effector-dependent neural activity.

### Effector versus colour activation across encoding, planning, and execution

#### Visual Target Encoding

During the *visual target encoding* phase, activity was observed within the left TP, left STG, and left MidFG. Activation of STG and MidFG is consistent with their established roles in spatial working memory (Geier et al., 2006) and spatial attention (Corbetta et al., 1998). MidFG, in particular, has been repeatedly implicated in the active maintenance of spatial location, though this lateralization is more commonly right-hemispheric (Owen et al., 1998; D’Esposito et al., 1998; Geier et al., 2006).

The left-lateralization of activity here be a result of participants preparing both reach and saccade responses before the action instruction. In that case, activity within these areas could be an early response for reach preparation with the right hand. In fact, Chen et al. (2014) similarly reported left MidFG activation during target encoding and their participants were aware of an upcoming required reach. The pattern of functional activity in these areas could also be related to the established left-lateralized network involved in language processing of action words (Boulenger et al., 2009). Thus, STG and MidFG activation likely reflect target location encoding, attention to a peripheral stimulus, early movement preparation, and language processing of the provided instructions.

The role of TP is less clear since this region is more associated with autobiographical memory and semantic-language processing, amongst other functions (Dupont, 2002). Its recruitment here may reflect its broader function as a multimodal convergence zone (Pascaul et al., 2015), supporting higher order integration of the spatial and contextual features of the target before upcoming movement.

#### Movement Planning

SACCADES. During saccade planning, activity was relatively limited, confined primarily to posterior parietal regions, including bilateral left SPL, and right PCu. There was also a small cluster within the left precentral gyrus. This pattern broadly agrees with previous studies isolating saccade planning activity (Astafiev et al., 2003; Beurze et al., 2009; Gallivan et al., 2011), although they reported more extensive frontal recruitment during this phase. The restrained activation here is consistent with preliminary findings from Cappadocia et al. (2018) and supports the interpretation that the saccade network does not become fully engaged until closer to the time of execution.

REACHES. As hypothesized, reach planning produced broader activation spanning occipital, parietal, and frontal cortices with peaks in the left SPL, left SFG, and left FP. This distributed pattern aligns with established findings and likely reflects the more elaborate processing required within the frontal cortex for upper limb movements (Astafiev et al., 2003; Gorbet et al., 2004; Beruze et al., 2007; 2009; Gertz and Fiehler, 2015; Cappadocia et al., 2017). The differential patterns observed here for saccade versus reach highlight the importance of separating planning from encoding and execution stages of movement.

#### Movement Execution

SACCADES. During saccade execution, strong bilateral engagement emerged across the bilateral SPL, bilateral precentral gyrus, left Occipital Fus., left PCu, and left FP. This pattern aligns well with prior neuroimaging work (Astafiev et al., 2003; Koyama et al., 2004; Simo et al., 2005; Filimon et al., 2009). The recruitment of Occipital Fus. within the fusiform gyrus is notable, as this region is not typically reported in saccade studies, though its recruitment has been document (Corbetta et al., 1998). Its engagement here may reflect the reactivation of a visual memory for the target’s location (Manelis et al., 2011). Alternatively, given the large extend of the cluster across much of the occipital cortex, activity here as a whole could represent visual processing as the eyes shift (Hagler et al., 2007; Levy et al., 2007).

REACHES. As hypothesized, reach execution produced broader cortical activation, with a left-lateralized distribution and peaks in the left postcentral gyrus and left precentral gyrus. These findings agree with the strong, contralateral somatosensory and primary motor involvement reported for reaching (Astafiev et al., 2003; Medendorp et al., 2005; Filimon et al., 2009; Cappadocia et al., 2017). While we did not observe any peaks within the well-characterizes regions of the posterior parietal and premotor cortices, the spatial pattern of our activation cluster confirmed the engagement of these regions. In addition, reach engaged the right lobule V, right lobule VIIIb, and vermis VIIIa of the cerebellum. The ipsilateral engagement within these lobules is consistent with somatotropic maps of upper limb representation in the cerebellum (Grodd et al., 2001) and their engagement during reach tasks (Nitschke et al., 2004).

### Overlapping Saccade and Reach activation

During *movement planning*, overlapping cortical activation for reach and saccades was confined to the same regions identified independently for saccade planning: bilateral SPL, right PCu, and left precentral gyrus. This pattern persisted into *movement execution*, where bilateral SPL and bilateral precentral gyrus, as well as the left PCu showed conjunctive activity. These findings agree with the hypothesis that shared overlap is present between these effectors and converges with a broad literature implicating the same parieto-frontal regions in both saccade and reach processing (Astafiev et al., 2003; Medendorp et al., 2005; Levy et al., 2007; Beurze et al., 2009; Filimon et al., 2009; Gallivan et al., 2011). The fact that shared activity was the same as the results found for saccade processing further confirms that reach and saccade recruit similar regions, with reaches engaging additional areas. Critically, the observed overlap here does not speak to the relative magnitude of activity for each effector, but it does offer an explanation for why dissociating effector-specific signals have proven to be difficult, especially when these networks are not directly contrasted with each other.

The shared role of spatial attention in these tasks must also be considered. It has been demonstrated that the saccade and spatial attention networks engage much of the same parieto-frontal regions (Corbetta et al., 1998; Astafiev et al., 2003). As both our reach and saccade conditions require spatial attention to remembered target location, our conjunction results may partly reflect the similar attentional demands required for both of these movements.

Overlapping activation was also identified in the cerebellum, encompassing hemispheric and vermal regions (including lobules VI, Crus I, and VIIIa, extending medially into vermis VI and VIIIa). Activity here did not coincide with peaks reported in either saccade or reach contrast independently. Still, this overlap is consistent with the established involvement of these regions across separate saccade and reach execution tasks (Nitschke et al., 2004; Diedrichsen et al., 2005; Errante and Fogassi, 2020; Yamamoto et al., 2025). Thus, the present study suggests that these regions might play similar supporting roles in both effectors.

### Effector-preferential activation across planning and execution

#### Movement Planning

SACCADES. Direct saccade and reach contrast during *movement planning* revealed limited saccade-preferential activity, with peaks in the bilateral Temporal Fus., right LOC, and left lobule I-IV in the cerebellum. The bilateral Temporal Fus. activation is unexpected given that this region is mostly associated with processes such as face recognition (Haxby et al., 1994), rather than oculomotor control. However, it may be involved spatial memory reactivation during saccade preparation (Manelis et al., 2011). More broadly, the lack of substantial dorsal occipital-parietal-frontal engagement is consistent with our hypothesis and previous findings showing exclusively reach-preferential activity during this phase (Cappadocia et al., 2018). This was also reinforced by ROI analyses, which revealed no regions, except the right mIPS, had saccade-related activity greater than the colour condition.

Saccade-related activity is typically observed in more posterior cerebellar regions (Nitschke et al., 2004; Simo et al., 2005; Yamamoto et al., 2005) while reach-related activity has been found in more anterior areas (Nitschke et al., 2004; Errante & Fogassi, 2020), which aligns with what we found during movement execution. Nevertheless, Alverez et al. (2010) reported increased activity for predictable versus random saccades within more anterior areas. Although target location was known for both effectors at this stage, activity in lobule I-IV may be related to the predictive processing of saccadic eye movements.

REACHES. In contrast, reach-preferential activity was broad, with peaks in the left SPL and left LOC. Moreover, the right aCing, bilateral DLPFC, MedFG, bilateral MidFG, bilateral PMd, bilateral M1, left AG, left mIPS, and bilateral SPOC all revealed significantly greater activity favoring this effector. These findings once again confirm that reach-planning activity is broader and stronger across the visuomotor network, at least earlier on during the sensorimotor stages (Astafiev et al., 2003; Medendorp et al., 2005; Beurze et al., 2009; Cappadocia et al., 2018).

Particularly for the PPC, previous studies have implicated the reach-specific role of SPOC in forming internal representations of spatial goals during reach planning (Fernandez-Ruiz et al., 2007), as well as mIPS and AG in integrating visual and proprioceptive information about hand position to form a reach vector (Vesia et al., 2010; Vesia and Crawford, 2012). Thus, our findings reflect the network involved in transforming sensory information into reach plans.

#### Movement Execution

SACCADES. The effector-preferential activity became most pronounced during *movement execution*. Saccade-preferential cortical engagement was observed across frontal, lateral posterior parietal, and occipital regions. Peaks were reported in the left Occipital Fus., left LG, left LOC, and left AG/SMG. ROI analyses revealed significantly greater saccade activity in left mIPS and right V1. Saccade preference within the occipital cortex is well-established (Hagler et al., 2007; Levy et al., 2007; Filimon et al., 2009) and can be attributed to the motion of the retinal image and visual input as the eyes shift fixations (Hagler et al., 2007; Levy et al., 2007; Filimon et al., 2009). Activation of AG and SMG during saccade tasks is well established (Simo et al., 2005; Schraa-Tam et al., 2009; Yamamoto et al., 2025), though whether these regions are preferentially active for saccades compared to reaches is less clear.

Previous fMRI work has shown reach-preference in AG (Filimon et al., 2009), while TMS evidence suggests this region plays a causal role in both reach and saccade movements (Vesia et al., 2010). Our cue-separated paradigm suggests that saccades preferentially activate these lateral PPC areas during movement execution. This would be consistent with the medial-lateral gradient that has been proposed along the IPS (Levy et al., 2007; Filimon et al., 2009) and extend this organization to the broader PPC. The greater activation for saccades in left mIPS also conflicts with previous reports of reach-preference (Filimon et al., 2009). During visuomotor preparation, left mIPS has been shown to decode spatial information rather than effector-type (Gallivan et al., 2011). The greater saccade activity in mIPS during execution may therefore reflect the later spatial processing of the saccade network. Despite our hypothesis, there were no peaks in the posterior vermis, although a small cluster was observed with the left lobule VIIb, which is consistent with previous findings (Guillaume et al., 2018). The lack of preferential activity within the oculomotor vermis (VI-VII) could be due to arm movements also recruiting similar areas (Nitschke et al., 2004). Previous reports comparing eye versus hand tracking also led to no significant eye-related activation relative to hand movements (Miall et al., 2002).

REACHES. Reach-preferential activity was observed in the left postcentral gyrus and left LOC, as well as right lobule V and right lobule VIIIb. The ROI analysis revealed significantly greater reach-related activity in the right aCing, MedFG, left MidFG, bilateral PMd, left M1 and bilateral SPOC, which have been previously established (Astafiev et al., 2003; Filimon et al., 2009; Cappadocia et al., 2018). Reach-preferential activity in the left postcentral gyrus, as well as the activation extension into medial PPC regions, is consistent with our hypothesis (Levy et al., 2007; Filimon et al., 2009; Cappadocia et a., 2018), although the absence of peaks within discrete medial PPC areas precludes strong region-specific claims.

Finally, the reach-preferential peak in the right lobule V and right lobule VIIIb once again confirms the established cerebellar representation of upper limb movements in the anterior and posterior lobes (Grodd et al., 2001), further validating the activation of these areas in prior reach tasks (Nitschke et al., 2004; Errante and Fogassi, 2020). Since saccade and reach activity has not previously been directly contrasted in the same manner shown here, our findings might also newly indicate effector-specific roles of the right lobule V and lobule VIIIb in reach execution.

### Multivoxel patterns

The multivariate findings revealed a more restricted distribution of significant saccade-reach spatial activity differences during planning compared with execution. Following removal of mean regional activation, significant distances were observed in only six ROIs during planning, including bilateral FEF, right PMd, right mIPS, right aCing, and right cerebellar lobule VI. This more restricted distribution may reflect the common sensorimotor processes involved in planning both eye and hand movements, including remembering the target location and transforming visual information into a motor goal. These shared demands may contribute to similar activity patterns within overlapping parieto-frontal regions. Nevertheless, the significant planning-related differences observed in bilateral FEF, right PMd, and right mIPS are consistent with previous multivoxel findings demonstrating effector-dependent preparatory activity within parieto-frontal regions, particularly PMd/FEF and right mIPS (Gallivan et al., 2011).

In contrast, movement execution requires distinct oculomotor and limb motor commands, accompanied by different sensory consequences and feedback processes. These differences may contribute to the more widespread spatial differentiation observed during execution, where the same 24 of 28 ROIs exhibited significant distances both before and after removal of mean regional activation. In particular, bilateral S1 showed significant differences during execution but not planning after removal of mean regional activation, consistent with the greater involvement of somatosensory processing during limb movement. Similarly, significant differences across bilateral cerebellar lobules VI and VIIIb and vermis VI may reflect the distinct demands of predicting, monitoring, and adjusting eye and hand movements.

### Multivariate vs. univariate findings

Two findings are particularly noteworthy. First, right M1 exhibited significant demeaned saccade-reach distances during execution, despite showing no significant differences in mean activation between effectors in the univariate ROI analyses. Similarly, whole-brain univariate contrasts did not identify reach– or saccade-preferential activation within the right precentral gyrus. Thus, the multivariate findings suggest that effector-dependent differences, not detected by univariate analyses may be reflected in the spatial distribution of activity within regions such as right M1. Second, significant demeaned distances were observed across widespread cortical and cerebellar regions during execution, including regions showing overlapping activation for both effectors in the univariate analyses. In particular, while univariate analyses revealed overlapping cerebellar activation across bilateral cerebellar hemispheres and vermis, multivariate analyses identified significant demeaned distances bilaterally in lobules VI and VIIIb, as well as vermis VI. These findings suggest that overlapping regional recruitment does not necessarily imply equivalent spatial activity patterns and that effector-dependent differences may extend beyond those identified through conventional regional activation contrasts.

### Limitations

Several limitations should be considered when interpreting the present findings. First, limitations in eye-tracking accuracy during the MRI scan prevented us from directly confirming fixation throughout each trial. Nevertheless, extensive participant training, assessment of eye movements during Run 0, and online eye monitoring provided assurance of task compliance. Equipment malfunctions also prevented behavioural data collection for four participants; however, training before and communication throughout the experiment supported their ability to perform the tasks as instructed.

Second, several regions involved in saccade and reach control overlap with parieto-frontal networks associated with spatial attention (Astafiev et al., 2003). Although the control condition was designed to minimize general attentional contributions, these processes may have influenced activity during the effector-dependent phases.

Third, the sample included substantially more females than males, limiting the generalizability of our findings across sexes. Previous studies have reported sex-related differences in premotor and parietal activation during reaching and upper-limb movements (Gorbet and Sergio, 2007; Andrushko et al., 2023). Thus, the unequal sex distribution may have influenced the observed activation patterns, although these effects were not directly examined.

Finally, no direct statistical tests were performed to evaluate differences between phases. Similarly, although the univariate and multivariate analyses examined the same anatomically defined regions of interest, they differed in preprocessing and analytical approach, including spatial smoothing and contrasts relative to the colour-report condition in the univariate analyses. Consequently, comparisons across phases and analytical approaches were descriptive.

### Conclusion

Together, our findings confirm that temporally separating sensorimotor phases reveals differential temporal evolutions for saccade and reach, with characteristic cortical and cerebellar activity at specific time points. We confirmed that there is overlapping activity along the bilateral parieto-frontal network during saccade and reach planning and execution, highlighting a reason for the difficulties present in detecting effector-specific dissociations without direct contrasts.

When such contrasts were applied, a clearer picture emerged. During planning, saccade-preferential activity was minimal and confined to cerebellar and ventral regions, with the absence of dorsal parieto-frontal recruitment confirming the saccade network attains full engagement closer to execution. Conversely, reach-preferential planning activity was broadly present across the dorsal occipital-parietal regions. The most revealing findings appeared during the execution, where saccade-preferential activity in AG/SMG and reach-preferential activity extending to the medial PPC elaborates on the medial-lateral gradient previously proposed for these effectors.

Additionally, reach-preferential activity within the ipsilateral cerebellum sheds light on non-cortical areas exhibiting stronger engagement for reaches relative to saccades. Our ROI analysis further revealed that during movement execution, many cortical regions show significantly greater activity for reach compared to saccade, with the exception of right V1 and left mIPS. Importantly, multivariate analyses revealed reliably distinguishable saccade and reach activity patterns across voxels within overlapping cortical and cerebellar regions, particularly during execution. These differences persisted even after removing mean regional activation. Thus, these results confirm that effector-differential signals are present within the cortex and cerebellum, and they become most apparent when sensorimotor phases are separated in time and examining multivoxel activity patterns.

## Data and Code Availability

Data and code are available for requests that comply with federal data sharing policies, pending approval from the requesting researcher’s local ethics committee and agreement on how to share author credits on any resulting publication.

## CRediT

CR: Conceptualization, Data Curation, Formal Analysis, Investigation, Methodology, Visualization, Writing – Original Draft

KM: Data Curation, Formal Analysis, Investigation, Visualization, Writing – Original Draft JG: Methodology, Formal Analysis, Writing – Review & Editing

JDC: Conceptualization, Funding Acquisition, Project Administration, Supervision, Writing – Review & Editing

## Ethics Approval

Investigating cortical mechanisms underlying visuomotor control using functional Magnetic Resonance Imaging (fMRI) in humans. Certificate #: e2023-336

## Acknowledgements

The authors thank Saihong Sun and Dr. Xiaogang Yan for technical support, and Dr. Gaelle N. Luabeya and Dr. Lina Musa for analysis discussions.

